# Electrical Signaling in Cochlear Efferents is Driven by an Intrinsic Neuronal Oscillator

**DOI:** 10.1101/2022.06.27.497817

**Authors:** Hui Hong, Laurence O Trussell

## Abstract

Efferent neurons are believed to play essential roles in maintaining auditory function. The lateral olivocochlear (LOC) neurons, which project from the brainstem to the inner ear where they release multiple transmitters including peptides, catecholamines and acetylcholine, are the most numerous yet least understood elements of efferent control of the cochlea. Using *in vitro* calcium imaging and patch-clamp recordings, we found that LOC neurons in juvenile and young adult mice exhibited extremely slow waves of activity (~0.1 Hz). These seconds-long bursts of Na^+^ spikes were driven by an intrinsic oscillator dependent on L-type Ca^2+^ channels, and were not observed in prehearing mice, suggesting an age-dependent mechanism underlying the intrinsic oscillator. Using optogenetic approaches, we identified both ascending (cochlear nucleus) and descending (auditory cortex) sources of synaptic excitation, as well as the synaptic receptors used for such excitation. Additionally, we identified potent inhibition originating in the glycinergic medial nucleus of trapezoid body (MNTB). Conductance-clamp experiments revealed an unusual mechanism of electrical signaling in LOC neurons, in which synaptic excitation and inhibition served to switch on and off the intrinsically generated spike burst mechanism, allowing for prolonged periods of activity or silence controlled by brief synaptic events. Protracted bursts of action potentials may be essential for effective exocytosis of the diverse transmitters released by LOC fibers in the cochlea.

**Significance Statement:** The lateral olivocochlear (LOC) neurons, being the most abundant auditory efferent control of the ear, remained largely unexplored. Here we reported that LOC neurons displayed patterned electrical activity at an unusually slow pace (~0.1 Hz), mediated by a calcium-dependent intrinsic oscillator. This is surprising given the speed and precision were believed to be the currency of signaling in the lower auditory system. Optogenetic experiments determined the glutamatergic and glycinergic sources of synaptic inputs to these neurons, while conductance-clamp experiments revealed that synaptic activity acts like switches for turning on or off prolonged spike activity driven by the intrinsic oscillator. This extended spike activity may be essential for effective exocytosis of the diverse transmitters released by LOC fibers in the cochlea.

## Introduction

The auditory efferent system, formed by brainstem neurons that project to the inner ear, adjusts the sensitivity of hearing in different sensory environments (for review see 1, 2). There are two major types of auditory efferent neurons, the medial and the lateral olivocochlear systems (MOC and LOC, respectively). Cholinergic MOC neurons form a quick reflex pathway to the cochlea, so that sound-evoked release of acetylcholine from efferent terminals reduces cochlear gain by hyperpolarizing outer hair cells (3, 4). Underlying this function are specialized biophysical properties of MOC neurons that allow them to reliably encode sound intensity (5). Recent evidence also suggests that MOC neurons are involved in the generation/progression of tinnitus in patients with hearing loss (6, 7).

By contrast, our knowledge about the LOC system is limited, which is surprising given that LOC neurons outnumber MOC neurons 3:1 (8, 9). LOC neurons are located in the lateral superior olive (LSO) of the superior olivary complex (SOC, Figure 1A). Their axons terminate onto the dendrites of spiral ganglion neurons that form the auditory nerve and may release a remarkably diverse cohort of neurotransmitters and neuromodulators in the cochlea, including acetylcholine, dopamine, GABA, calcitonin gene-related peptide (CGRP), urocortin, and opioid peptides (1, 10). For decades, LOC neurons have been thought to play a role in auditory disorders, including noise-induced hearing loss, tinnitus and hyperacusis (11–14). However, due to the small caliber of LOC axons, no recordings exist of their activity *in vivo*. Moreover, previous studies *in vitro* were conducted either in young animals with immature hearing, or in older animals without a biomarker to distinguish LOC neurons from other cell types (15–17). Thus, there is little information about the function of these neurons, hindering our understanding about the control of the auditory system and its pathology.

**Figure 1.**
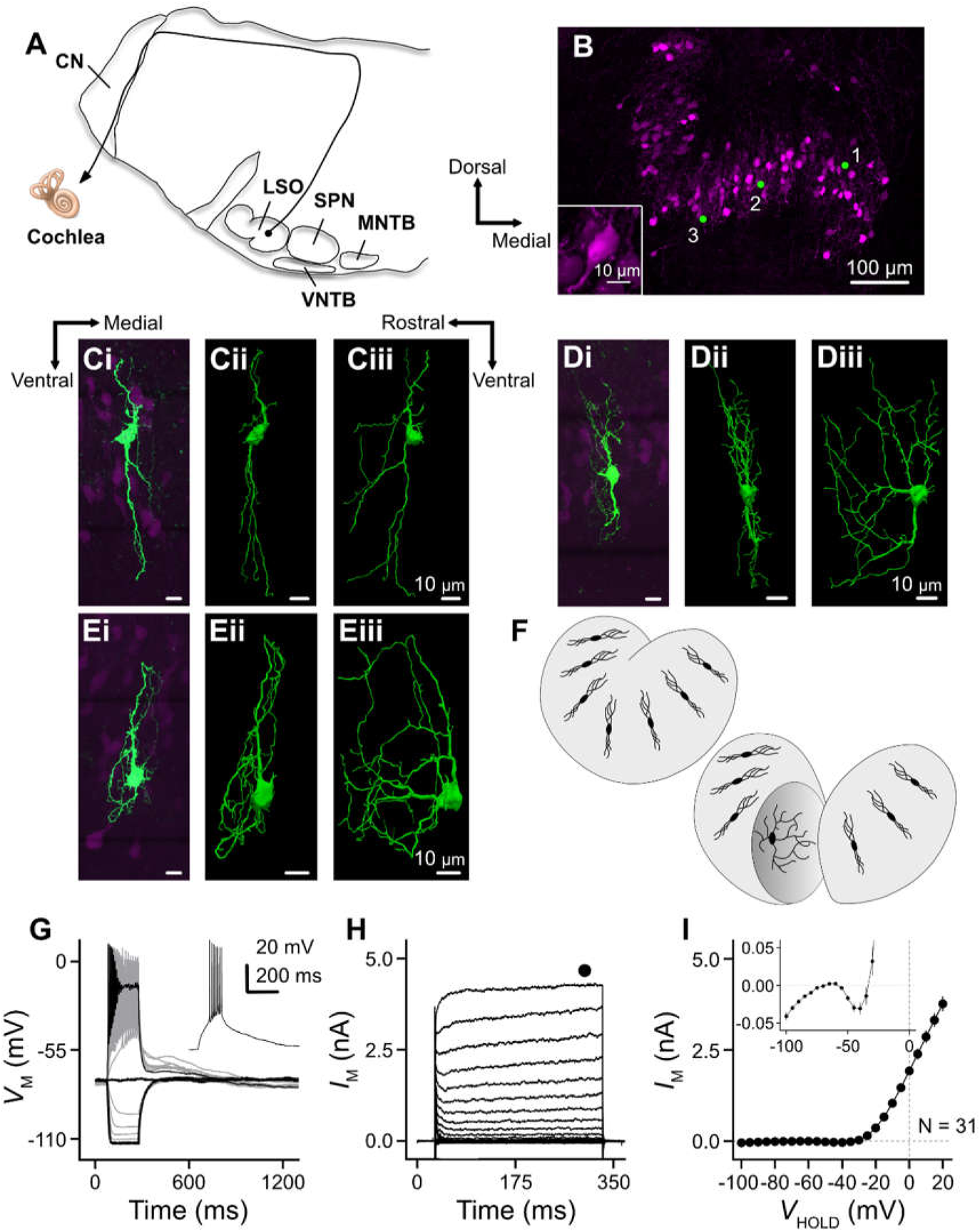
Morphological and biophysical properties of lateral olivocochlear (LOC) neurons in juvenile and young adult mice. (A) Schematic figure showing the coronal section of auditory brainstem containing cochlear nucleus (CN) and superior olivary complex (SOC), along with the axon projection from an LOC neuron to the cochlea. SOC contains nuclei such as lateral superior olive (LSO), superior paraolivary nucleus (SPN), medial and ventral nucleus of trapezoid body (MNTB and VNTB, respectively). (B) Coronal brainstem slice from a ChAT-Cre/tdTomato mouse showing the expression of tdTomato in the cholinergic LOC neurons. Numbers represent the location of three neurons shown in (C). Inset shows an enlargement of an LOC neuron. Dorsal to the top and medial to the right. (C-E) Ci-Ei: Representative confocal images of three LOC neurons filled with 1% biocytin. Cii-Eii: The most restricted view of dendritic field after reconstruction. Ciii-Eiii: The most extensive view of dendritic field after reconstruction. (C-E) share the orientation bars. (F) Schematic illustration showing the bipolar appearance of LOC neurons in the coronal plane (upper) but a more expansive distribution of dendrites in the parasagittal plane (lower). (F) Representative voltage traces of an LOC neuron in response to current injections from −100 to 200 pA in steps of 20 pA. Thickened traces represent voltage response to 200, 0 and −100 pA current injections. Inset shows the voltage response to 20 pA current injection. (G) Representative current traces of an LOC neuron in response to V_H_ from −100 to +20 mV in steps of 5 mV. Filled circle denotes the measurement window of steady-state current shown in (F). (H) Population data showing the I-V relation of steady-state current. Inset shows an enlargement of current at V_H_ between −100 and 0 mV. N, the number of neurons. Error bars, SEM.

Here, we reveal the basis for electrical activity in LOC neurons of juvenile and young adult mice. By applying calcium imaging and patch-clamp techniques, we observed a novel, infra-slow (~0.1 Hz) pattern of intrinsic activity in LOC neurons, characterized by low-frequency firing of seconds-long bursts of Na^+^ channel-dependent action potentials (APs) driven by slow, Ca^2+^ channel-dependent voltage waves. Using optogenetics, we determined that LOC neurons receive ascending excitatory inputs from T-stellate cells in the cochlear nucleus (CN, Figure 1A), descending excitatory inputs from auditory cortex, and inhibition from the medial nucleus of trapezoid body (MNTB) of the SOC (Figure 1A). Unlike convention neuronal excitation, even a short period of synaptic activity acted to turn on and off the prolonged Ca^2+^-dependent spike trains, like a light switch. Thus, we propose that the output of the LOC system is a synaptically-triggered extended spike burst that may be essential for the release of a variety of conventional transmitters and peptides in the cochlea.

## Results

### Morphological and biophysical properties of LOC neurons

We used transgenic mice of both sexes expressing Cre recombinase under the endogenous choline acetyltransferase promoter (ChAT-IRES-Cre) and crossed them with a tdTomato reporter line (Ai9, see Materials and Methods for details). These mice are referred to as ChAT-Cre/tdTomato. Thus, cholinergic LOC neurons in the LSO expressed the red fluorescent protein, tdTomato, and could be visualized in brainstem slices for electrophysiological recordings (Figure 1B). The distribution of LOC neurons displayed an “S” shape in coronal slices characteristic of the shape of the LSO. Filling LOC neurons with 1% biocytin further revealed that these neurons had small, ovoid cell bodies with bipolar dendrites extending across the dorsoventral axis of the LSO (Figure 1Ci-Ei). The approximate location of three labeled neurons within the LSO is indicated in Figure 1B. Similar to LSO principal cells, the long axes of LOC dendrites ran perpendicular to the curvature of the LSO, varying with tonotopic regions (18, 19). Individual neurons were reconstructed to reveal the dendritic distribution in three dimensions. While the raw images showed a bipolar morphology, the reconstructions revealed that the neurons were disc-shaped, projecting their dendrites along narrow bands into the rostro-caudal axis. Figures 1Cii-Eii illustrate reconstructions after small rotations (~10°) around the medial-lateral (coronal) plane width and revealed the narrowest extent of dendrites, while Figures 1Ciii-Eiii show reconstructions after further rotations of 90°, i.e., extending into the rostro-caudal (parasagittal) axis. For both image angles, a contour was drawn around the neurons to include their most distal processes. The area of each contour was calculated and showed that the dendritic fields were 2.55 ± 0.25 times (n = 9 neurons) more extensive in the parasagittal than coronal planes. This narrow projection is illustrated in the cartoon in Figure 1F and strongly suggests that LOC neurons receive narrowly-tuned synaptic signals. Upon break-in during whole-cell recording, the neurons were initially silent, allowing us to measure passive properties near −60 mV. In current-clamp mode, passive membrane properties of LOC neurons were characterized by injecting a −5-pA current (20). The exponential time course and amplitude of the voltage response yielded a time constant of 57.42 ± 4.05 ms, input resistance of 1.19 ± 0.09 GΩ, and membrane capacitance of 50.79 ± 2.81 pF (n = 33 neurons, also see Figure S1A). The extremely high resistance indicates that few ionic conductances were active near −60 mV. To study active properties, a series of currents with the amplitude from −100 pA to 200 pA were injected into LOC neurons with steps of 20 pA. Representative voltage responses are shown in Figure 1G. Because of their high input resistance, only 20 pA was needed to evoke APs (Figure 1G, inset), and 200 pA was sufficient to induce depolarization block (Figure 1G). Conductance increased in response to negative current injections, likely due to the activation of inwardly-rectifying potassium channels (Figure 1G). In whole-cell voltage-clamp mode, LOC neurons were held at voltages between −100 mV to +20 mV and current-voltage (I-V) relations were plotted (Figure 1H, 1I). At membrane voltage positive to −30 mV, outward current increased, presumably carried by high-voltage activated potassium channels. However, between −55 mV and −75 mV, currents hovered near 0 pA, underlying the exceptionally high input resistance of LOC neurons.

### Calcium imaging revealed infra-slow spontaneous activity in LOC neurons

During current-clamp experiments, spontaneous firing of LOC neurons emerged within 5 minutes after establishment of whole-cell mode, as described in detail below. As we were concerned that such activity might be a result of slight damage due to the recording pipette, we employed a non-invasive measure of activity using calcium imaging. AAV1-CAG-Flex-mRuby2-GSG-P2A-GCaMP6f-WPRE-pA (21) was injected into the LSO of ChAT-Cre mice at postnatal day (P) 21-23 (Figure 2A, see Materials and Methods). Brainstem slices were harvested 1-2 weeks after the surgery and imaged under two-photon microscopy. This viral construct caused cholinergic LOC neurons to be visualized by expression of red fluorescent protein mRuby while their electrical activity could be detected by the Ca^2+^ indicator GCaMP6f. Figure 2 B1-B2 shows six representative LOC neurons at two different time points. Fluorescent signals of GCaMP6f for each neuron were first normalized to their corresponding mRuby signal. This F_G_/F_R_ was then scaled to a range between 0% and 100% (“Scaled F_G_/F_R_”, Figure 2C and S2A). The mean and standard deviation (SD) of the baseline fluorescence were calculated (Figure S2B, see Materials and Methods) and LOC neurons were categorized as spontaneously active if their scaled F_G_/F_R_ exceeded 4 SDs from the averaged baseline (Figure S2A). Oscillation frequency was measured based on the auto-correlation of scaled F_G_/F_R_ for each neuron (Figure S2C). By this analysis, 90% of LOC neurons we imaged showed spontaneous Ca^2+^ signals with a mean oscillation frequency of 0.110 ± 0.004 Hz (n = 170 oscillating neurons).

**Figure 2.**
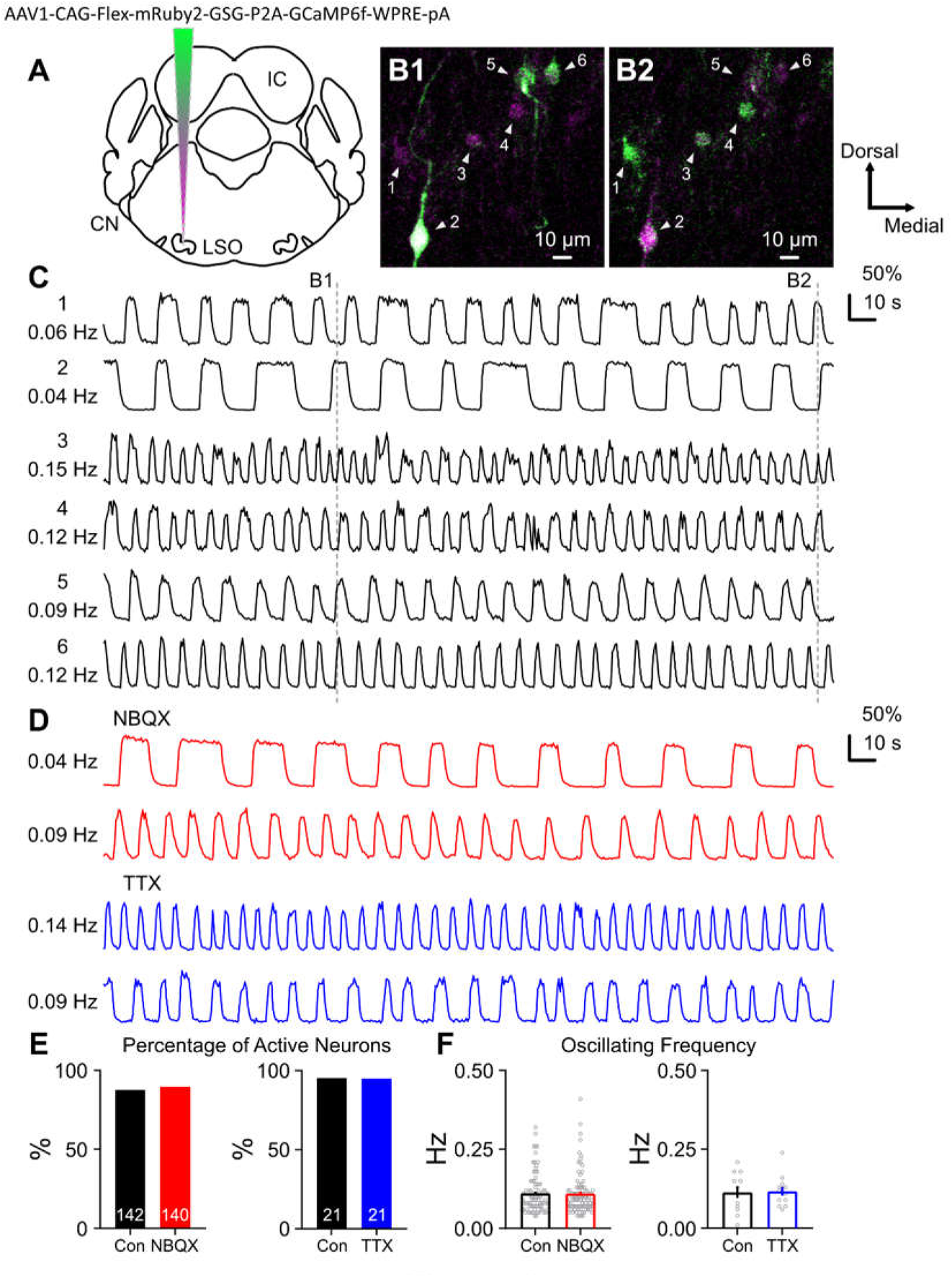
Calcium imaging revealed infra-slow spontaneous electrical activity in LOC neurons. (A) Schematic figure showing the injection of Cre-dependent virus AAV1-CAG-Flex-mRuby2-GSG-P2A-GCaMP6f-WPRE-pA into the LSO. (B1-B2) Two-photon scanning at two different time points. Magenta channel represents the expression of mRuby, while green channel (on and off) represents the activity of GCaMP6f. Arrowheads point to six representative LOC neurons, the scaled F_G_/F_R_ of which is shown in (C). (C) The scaled F_G_/F_R_ of six representative neurons plotted as a function of time. The oscillating frequency for each neuron is listed on the left. Two dash lines denote two time points, the real-time scanning of which is shown in (B1) and (B2). (D) Representative traces of scaled F_G_/F_R_ in response to bath application of NBQX (20 µM, red) or TTX (0.5 µM, blue). Note that two neurons in NBQX are Neuron 2 and 5 shown in (C), respectively. (E) Bar graph showing the percentage of active neurons in control (con), with bath application of NBQX or TTX. Numbers in the bars represent the number of neurons. (F) Population data showing the oscillating frequency in control (con), with bath application of NBQX or TTX. Error bars, SEM.

In order to determine if the Ca^2+^ signals were driven by synaptic activity, we applied NBQX (20 µM) to block AMPA and kainate receptors. NBQX did not block spontaneous Ca^2+^ signals (Figure 2D, red traces); the percentage of LOC neurons that were active was 87.8% in control and 89.6% with NBQX (Figure 2E, left panel). Oscillation frequency also did not change with drug (Figure 2F, left panel, *p* = 0.946; paired *t* test). Thus, spontaneous Ca^2+^ signaling of LOC neurons is not driven by spontaneous activity of glutamatergic inputs in the brain slice. Surprisingly, the Na^+^ channel antagonist TTX (0.5 µM) also failed to affect the Ca^2+^ signals (Figure 2D, blue traces; Figure 2E, right panel, control: 95.5%, TTX: 95.0%; Figure 2F, right panel, *p* = 0.818; paired *t* test), indicating that spontaneous Ca^2+^ signaling in LOC neurons is not dependent on the generation of Na^+^-dependent APs.

### Electrical activity in LOC neurons is dependent on L-type Ca^2+^ channels

We next asked whether the spontaneous activity in the electrophysiological recordings had properties similar to the spontaneous Ca^2+^ signals described above. Recordings were made in both cell-attached and whole-cell current-clamp mode, and the pattern of spike activity compared to the pattern observed with imaging. In cell-attached and whole-cell experiments, LOC neurons exhibited seconds-long trains of spikes, interrupted by lengthy silent periods (Figure 3A), not unlike the infra-slow pattern of activity seen with imaging. In a subset of experiments, gramicidin-perforated patch-clamp recordings were used instead of whole-cell recordings (n = 5 neurons). We observed a similar pattern of bursting activity and thus these data were pooled with whole-cell data. During the hyperpolarizing phase of the bursts, neurons often exhibited a voltage “shoulder” around −60 mV (Figure 3A, lower panel, inset, arrowhead), a membrane voltage at which near-zero current was present (see Figure 1I, inset), suggesting that cessation of bursts represents a transition among “bistable” membrane potentials. In this analysis, we distinguish between frequency of individual spikes and the frequency of spike bursts, that is, the rate at which bursts arise. Figure 3C illustrates the histogram of intervals between spikes within individual bursts (i.e., inter-spike interval, ISI). The peak occurs at an ISI of 40 ms (i.e., a spike frequency of 25 Hz). However, the frequency of bursts was not significantly different from the frequency of Ca^2+^ waves (Figure 3D, *p* = 0.278, ANOVA). Thus, the long burst of spikes and the long Ca^2+^ waves may be driven by similar mechanisms.

**Figure 3.**
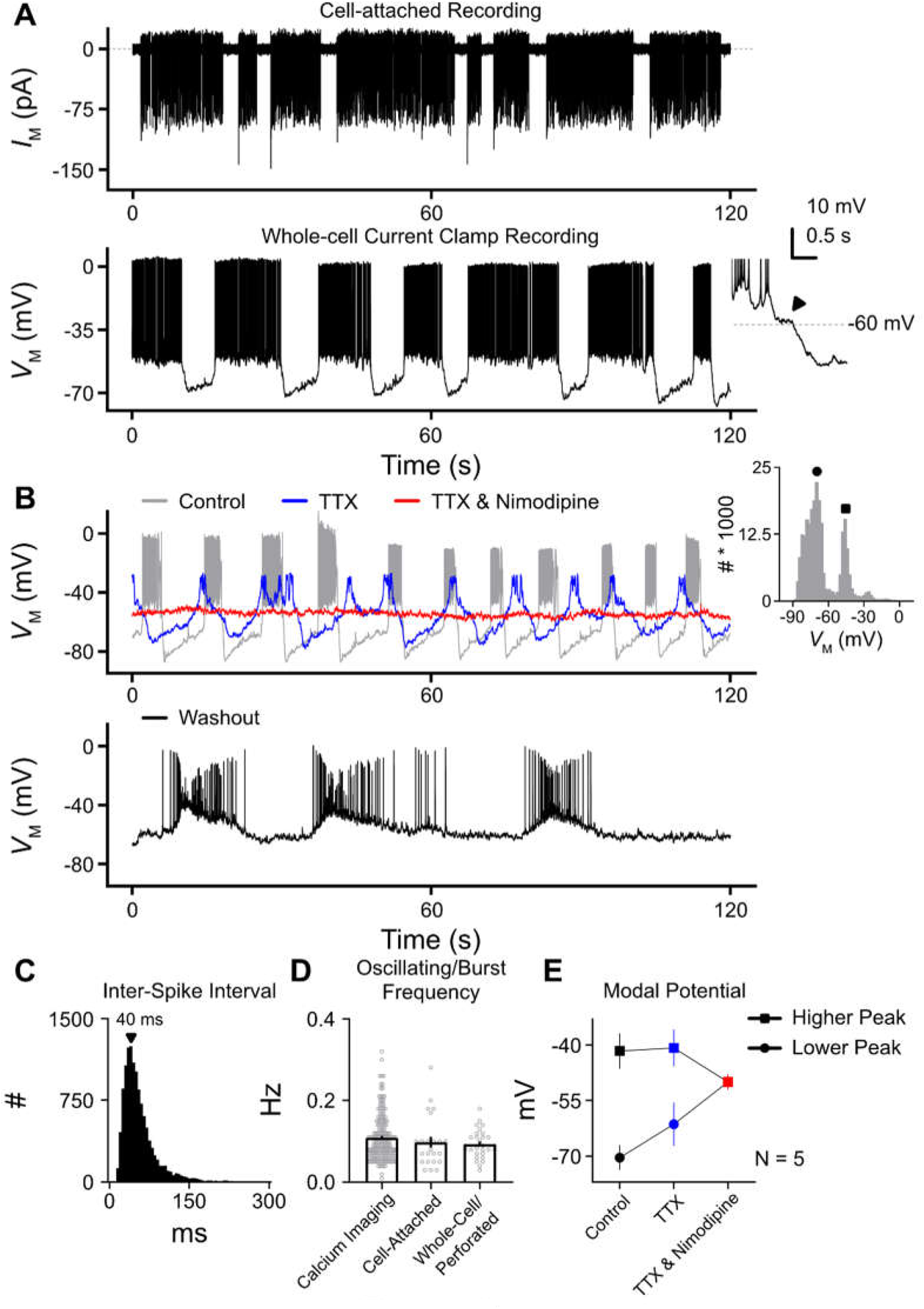
Infra-slow spontaneous electrical activity in LOC neurons is dependent on L-type calcium channels. (A) Representative traces showing the spontaneous activity of LOC neurons obtained using cell-attached recording (upper panel) or whole-cell current-clamp recording (lower panel). Inset shows the enlargement of cessation phase of a burst. Arrowhead points to the typical voltage “shoulder” that precedes the completion of the hyperpolarization after a burst. (B) Representative spontaneous activity of an LOC neuron in control, with bath application of TTX (0.5 µM) and nimodipine (3 µM, upper panel) and after washout of these two drugs (lower panel). Inset shows the histogram of membrane voltage in control. The lower and higher peak (i.e., modal potentials), denoted by circle and square, respectively, is plotted in (E). No bias current was applied to this LOC neuron. (C) The histogram of inter-spike intervals (ISI) within the bursts. Arrowhead points to the peak at 40 ms. (E) Population data showing the oscillating or burst frequency obtained using four different techniques. Note that the data from whole-cell and perforated patch-clamp recordings are pooled. (F) Population data showing the modal potentials acquired from the histogram of membrane voltage (see B, inset) and plotted across three different conditions. N, the number of neurons. Error bars in (D) and (E), SEM.

To probe further the basis for these slow oscillations, we applied 0.5 µM TTX to the bath. While, as expected, fast AP activity was eliminated, we still observed prominent membrane potential oscillations (Figure 3B, upper panel, blue trace), consistent with the results from imaging in which TTX did not alter Ca^2+^ oscillations. Next, we washed in nimodipine (3 µM), an L-type Ca^2+^ channel antagonist, in addition to TTX. Nimodipine eliminated all voltage oscillations, leaving the membrane in a partially depolarized state (Figure 3B, upper panel, red trace). Burst firing returned when the two drugs were washed out (Figure 3B, lower panel). A histogram of membrane voltage was plotted for these three conditions, and an example from the control is shown in the inset of Figure 3B. For this neuron, the bimodal distribution contains two peaks that occur at −70 mV and −40 mV. Plots were made of the membrane voltage at the histogram peaks (i.e., modal potentials) across the three different conditions (Figure 3E). The difference between the peaks became smaller with TTX, and the two peaks merged into one upon addition of nimodipine, as oscillations were abolished. This reduction in the peak and trough of the oscillation by TTX presumably results from reduced activation of the repolarizing K^+^ current which turns on positive to −20 mV (Figure 1I). Taken together, an L-type Ca^2+^ channel-dependent intrinsic oscillator produces slow periodic changes in voltage that in turn trigger long burst firing of APs.

### Intrinsic properties of LOC neurons in prehearing mice

Previous studies of LOC neurons focused on young, prehearing rodents (15, 16). To explore the age-dependent differences in passive and active biophysical properties, we also recorded LOC neurons at P9-11, prior to hearing onset in mice (P12, ref 22). In current clamp, only 44% of LOC neurons from prehearing mice showed spontaneous activity, as compared to 100% for the older mice. Moreover, when active, prehearing LOC neurons fired tonically rather than in a burst pattern (Figure S1B, left panel). The histogram of ISIs peaked at 110-150 ms, corresponding to spike frequency of 6.7-9.1 Hz (Figure S1B, right panel), much slower than the spike rate in older mice (25 Hz). Immediately after whole-cell configuration, spontaneously active neurons were temporarily silent, which allowed us to measure their passive membrane properties. Neurons at these ages (including both spontaneously active and inactive neurons) had significantly smaller input resistances (0.75 ± 0.06 GΩ; Figure S1A, *p* = 0.001, unpaired *t* test) but larger membrane capacitance (86.87 ± 9.37 pF; *p* < 0.0001, unpaired *t* test). The time constant of the membrane (58.50 ± 3.49 ms) did not show a difference with age (*p* = 0.860, unpaired *t* test). Voltage responses of prehearing LOC neurons to current injections from −100 pA to 200 pA resembled those from older neurons (Figure S1C), and overall our data from younger neurons are similar to those reported in previous studies (15, 16). Taken together, with the onset of hearing, LOC neurons gain the capacity for burst firing, likely through strengthening an intrinsic voltage oscillator mechanism.

### LOC neurons receive ascending excitatory inputs from the CN

The prominent intrinsic activity we described above must be guided by synaptic activity. However, little is known about what neurons or brain regions provide input to LOC neurons. Because LOC neurons may receive ascending excitatory inputs from the CN (16, 23), a Cre-independent virus (24) that carries a sequence for channelrhodopsin (ChR2) fused to the fluorophore Venus was injected into the CN of ChAT-Cre/tdTomato mice (Figure 4A, see Materials and Methods), with the expectation that terminals of CN neurons would express ChR2. The arrowhead in Figure 4B points to the infected CN expressing Venus. 1-2 weeks after the surgery, voltage-clamp experiments were conducted on LOC neurons ipsilateral to the injection site. Trains of light pulses (5, 10 and 20 Hz) elicited robust excitatory postsynaptic currents (EPSCs) in LOC neurons (Figure 4C and 4D, V_H_ (holding voltage) = −70 mV). These EPSCs were blocked by GYKI-53655 (50 µM), an AMPA receptor antagonist (Figure 4C) and their decay time constant (tau) was 4.27 ± 0.19 ms (Figure 4C, inset; n = 22 neurons). The reduction in the absolute amplitude of EPSCs along the pulse train (Figure 4D) was partially due to short-term synaptic depression as it was also observed when a stimulating electrode was used to excite input fibers to LOC neurons (Figure 4E and 4F). Moreover, we previously showed that ChR2 expression in projection neurons in CN enables reliable presynaptic AP activation at this frequency (5). However, it is possible that desensitization of ChR2 itself may also contribute to this decline (25).

**Figure 4.**
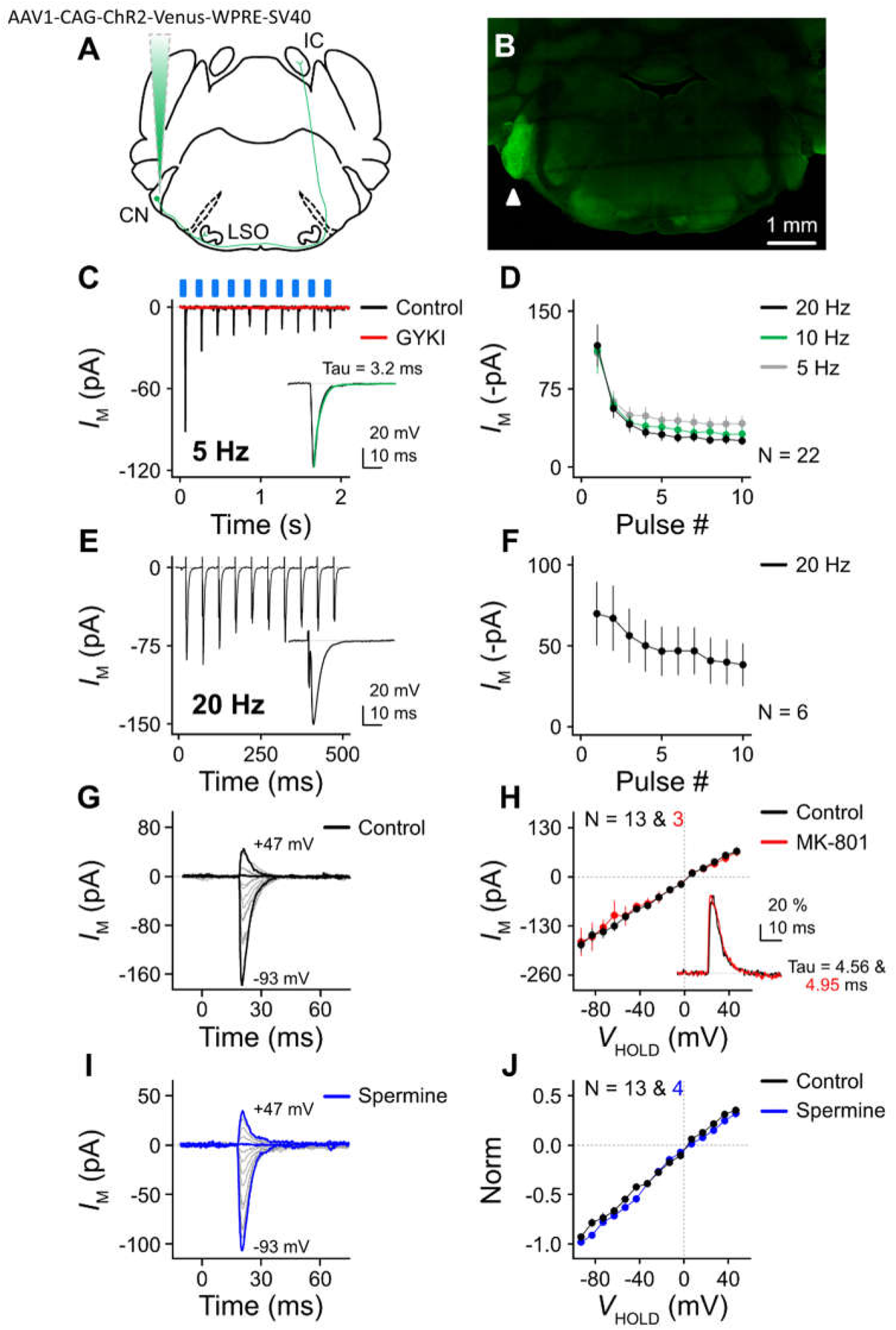
LOC neurons receive ascending excitatory inputs from the cochlear nucleus (CN) (A) Schematic figure showing the injection of anterograde-transported, Cre-independent virus AAV1-CAG-ChR2-Venus-WPRE-SV40 into the CN. (B) Acute brainstem slice with the arrowhead pointing to the infected CN. (C) Representative current traces of an LOC neuron in response to a train of light flashes at 5 Hz, in control and with bath application of GYKI-53655 (50 µM). Blue rectangles denote the occurrence of light stimulation. Inset shows the enlargement of the first EPSC. A single exponential fit to the decay phase (green curve) renders a decay time constant (tau) of 3.2 ms. (D) Population data showing the absolute amplitude of EPSCs as a function of light stimulus number. (E) Representative current traces of an LOC neuron in response to a train of electrical simulation at 20 Hz. Inset shows the enlargement of the first EPSC. (F) Population data showing the absolute amplitude of EPSCs as a function of electrical stimulus number. (G) Representative current traces of an LOC neuron in response to single light stimulation at V_H_ from −93 to +47 mV, in steps of 10 mV. (H) Population data showing the I-V relation of EPSC in control and with bath application of MK-801 (10 µM). Inset shows the overlaid, normalized EPSCs at +47 mV under these two conditions. (I) Representative current traces of an LOC neuron in response to single light stimulations at the V_H_ from −93 to +47 mV, in steps of 10 mV. Recordings were made with 0.1 mM spermine in the internal solution. Thickened traces in (G) and (I) represent V_H_ at −93, +7 and +47 mV. (J) Population data showing the normalized I-V relation of EPSC in control and with spermine in the recording pipette. N in (D), (F), (H) and (J), the number of neurons. Error bars in (D), (F), (H) and (J), SEM.

EPSCs were evoked at different holding potentials (Figure 4G) in order to construct I-V relations. While the AMPA receptor antagonist GYKI-53655 blocked all of the EPSCs at −70 mV, it remained possible that NMDA receptors could contribute to the EPSC at more positive potentials due to relief of Mg^2+^ block of the NMDA receptor channel. However, I-V relations were linear (Figure 4H), and the NMDA receptor antagonist MK-801 (10 µM) did not alter the shape of the EPSC even at positive potentials (Figure 4H, inset). Indeed, the average decay tau at the V_H_ between +17 mV and +47 mV was similar across the two conditions (control: 5.23 ± 0.14 ms; MK-801: 5.00 ± 0.30 ms; *p* = 0.622, paired *t* test, n = 3 neurons), again indicating the absence of NMDA receptors at these synapses.

The I-V relations of EPSCs exhibited little inward rectification, suggesting Ca^2+^-impermeable GluA2-containing AMPA receptors in LOC neurons (Figure 4H, ref 26). The slope of this I-V curve represents the excitatory conductance elicited by the fibers from CN, and was 1.786 nS. Since inward rectification of Ca^2+^-permeable AMPA receptors arises from channel block by intracellular spermine, and because spermine might be dialyzed out of the neuron during recording, we added 0.1 mM spermine to the recording pipette. The resulting EPSCs and their I-V curve still lacked inward rectification with spermine (Figure 4I and 4J). In summary, LOC neurons receive ascending excitatory inputs from the CN, mediated by Ca^2+^-impermeable GluA2-containing AMPA receptors but not by NMDA receptors. This property is in contrast to MOC neurons, which express Ca^2+^-permeable GluA2-lacking AMPA receptors that enable ultrafast synaptic transmission (5).

### Ascending excitatory input from T-stellate cells

Because T-stellate cells in the ventral CN send axon collaterals to LSO, they are a potential source of excitatory inputs to LOC neurons (27). T-stellate cells also project to contralateral inferior colliculus (IC), while the other major cell types in the ventral CN, such as bushy cells, do not. Therefore, to label T-stellate cells, we injected a retrograde-transported virus (28) into the IC (Figure 5A and 5B1). As illustrated in Figure 5B2, this virus expressed ChR2 along with GFP in the IC-projecting T-stellate cells located in the contralateral CN; if these T-stellate cells form collaterals to the ipsilateral LOC, we should be able to evoke EPSCs with light stimulation. Indeed, robust EPSCs were evoked in LOC neurons contralateral to the injected IC and exhibited a pattern of short-term synaptic depression similar to that seen with CN injection (Figure 5C, left panel, and 5D, V_H_ = −70 mV). Since the “retrograde” virus might also label local IC neurons and its ChR2 transported in an anterograde fashion, we injected a verified “anterograde” virus (mentioned above) in the IC to test if some IC somata might project to either ipsi- or contralateral LOC neurons. However, in recordings from 15 neurons (7 ipsilateral and 8 contralateral neurons to the injection site), no light evoked EPSCs were observed, indicating that the inputs we observed following retrograde virus injection are likely from T-stellate cells. Like the EPSCs evoked after viral injection into CN, the EPSCs evoked after retrograde virus IC injection exhibited no inward rectification (Figure 5F and 5G), indicating that IC-projecting T-stellate cells form ascending excitatory inputs to LOC neurons mediated by Ca^2+^-impermeable GluA2-containing AMPA receptors. However, although all the fast EPSCs were blocked by GYKI-53655, in 14 of 16 cells there remained a small, GYKI-resistant EPSC with much slower kinetics (Figure 5C, middle panel; decay time constant = 304.7 ± 43.6 ms). The amplitude of this slow EPSC was frequency-dependent, becoming more pronounced upon higher-frequency stimulation (Figure 5E, *p* = 0.0025, repeated-measure ANOVA). This current (recorded in the presence of GYKI-53655) was blocked by the AMPA/kainate receptor antagonist NBQX (10-20 µM), indicating that it was mediated by kainate receptors (Figure 5C, right panel). The kainate receptor-dependent input probably did not originate from the CN or IC, because injection of anterograde AAV to the CN did not lead to kainate receptor-dependent EPSCs, and as noted above, injection of anterograde AAV to the IC did not result in EPSCs in the contralateral LOC, consistent with previous histological reports that IC does not project to LSO (29, 30). Thus, there is an additional source of input to LOC neurons which sends collaterals to IC, and was infected by our retrograde virus. In the next section, we show that this additional source is the auditory cortex.

**Figure 5.**
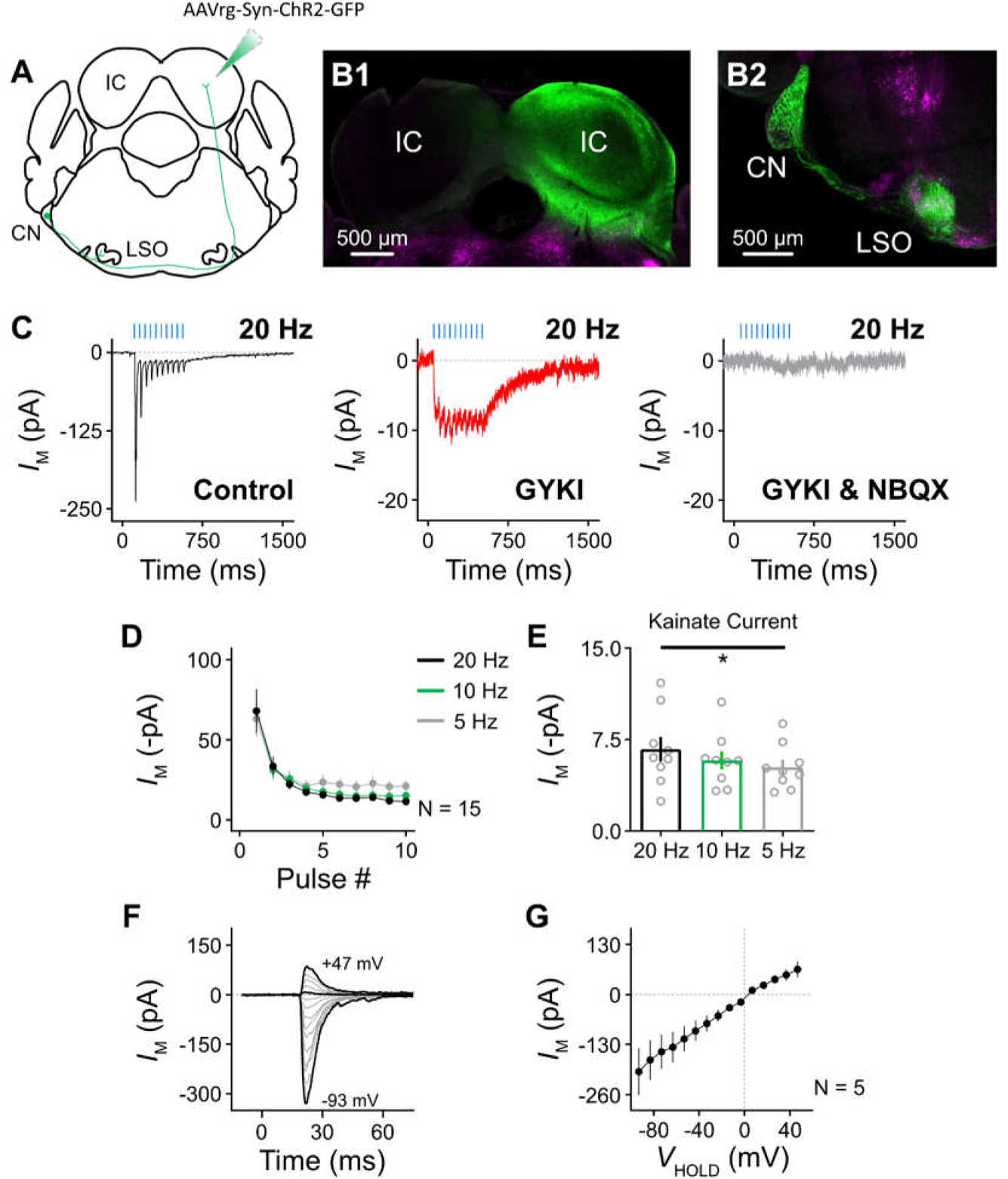
T-stellate cells provide ascending excitatory inputs to LOC neurons. (A) Schematic figure showing the injection of retrograde-transported, Cre-independent virus AAVrg-Syn-ChR2-GFP into the inferior colliculus (IC). (B1-B2) Acute brainstem slice showing the infected IC (B1) and the retrograde-labeled T-stellate cells in the contralateral CN sending fibers into the LSO. Magenta channel represents the expression of tdTomato, while green channel represents the expression of GFP. (C) Representative current traces of an LOC neuron in response to light stimulation of 20 Hz in control, with GYKI-53655 (50 µM), and with GYKI-53655 and NBQX (10-20 µM). Blue lines denote the occurrence of light stimulation. (D) Population data showing the absolute amplitude of EPSCs as a function of light stimulus number. (E) Population data showing the maximal kainate receptor-dependent current in response to light stimulation at different frequencies. * = *p* < 0.05 (repeated-measure ANOVA). (F) Representative current traces of an LOC neuron in response to single light stimulations at the V_H_ from −93 to +47 mV, in steps of 10 mV. Thickened traces represent V_H_ at −93, +7 and +47 mV. (G) Population data showing the I-V relation of EPSC. N, the number of neurons. Error bars in (D), (E) and (G), SEM.

### LOC neurons receive descending excitatory inputs from the auditory cortex

Previous histological studies showed that neurons in the auditory cortex send axons to both the IC and SOC (31, 32). As auditory cortex is thus a candidate source of descending input to LOC neurons, we injected the same ChR2-expressing anterograde virus into the auditory cortex (Figure 6A). This injection resulted in Venus labeling of fibers in the medial geniculate body ipsilateral to the injection site, one of the multiple targets of descending cortical fibers (Figure 6B, arrowhead). Additionally, sparse expression of Venus-positive boutons was observed in the contralateral auditory cortex (not shown). Sparse cortical fibers were also observed in the LSO, consistent with previous studies (Figure 6C, arrowheads, ref 31, 33). Importantly, some of these cortical fibers are in close proximity to tdTomato-labeled LOC processes (Figure 6C, inset). In line with these anatomical results, light flashes evoked EPSCs in LOC neurons, and a kainate-receptor component was observed in 2 out of 7 recorded neurons with stimulus-evoked EPSCs (Figure 6D, V_H_ = −70 mV). The absolute amplitude of fast EPSCs (i.e., AMPA receptor-dependent current) is plotted in Figure 6E and showed a pattern of short-term depression. I-V relations of the EPSCs were linear, again indicating the presence of GluA2 subunits (Figure 6F and 6G). MK-801 (10 µM) did not change the linearity of the I-V curve or alter the shape of EPSCs at positive potentials, indicating the absence of NMDA receptors at cortical-LOC synapses (Figure 6G and 6G, inset; average decay tau between +17 mV and +47 mV, control: 6.02 ± 0.89 ms; MK-801: 6.30 ± 0.78 ms; *p* = 0.486, paired *t* test, n = 4 neurons). In summary, LOC neurons receive descending inputs from the auditory cortex, the synapses of which employ both Ca^2+^-impermeable GluA2-containing AMPA receptors and in some cases, kainate receptors.

**Figure 6.**
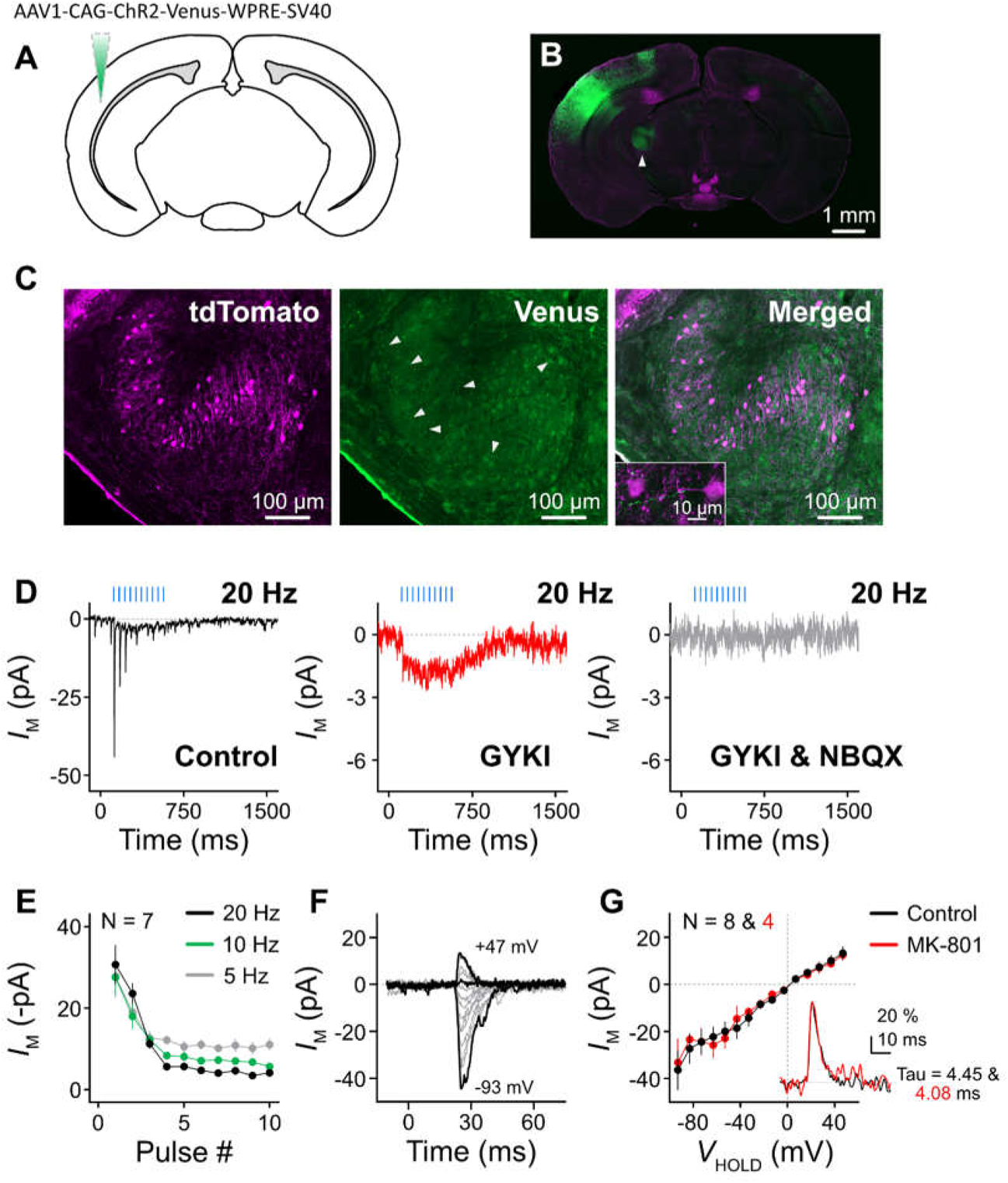
LOC neurons receive descending excitatory inputs from the auditory cortex. (A) Schematic figure showing the injection of anterograde-transported, Cre-independent virus AAV1-CAG-ChR2-Venus-WPRE-SV40 into the auditory cortex. (B) Brain slice showing the infected auditory cortex. Arrowhead points to the medial geniculate body. Magenta channel represents the expression of tdTomato, while green channel represents the expression of Venus. (C) Confocal imaging showing the expression of tdTomato (magenta) and Venus (green) in the LSO ipsilateral to the cortical injection site. Arrowheads point to the Venus-positive cortical fibers. Inset is the high-magnification confocal image showing the Venus-positive cortical fibers in close proximity to tdTomato-positive (i.e., LOC) neurons and processes. (D) Representative current traces of an LOC neuron in response to light stimulation of 20 Hz in control, with GYKI-53655 (50 µM), and with GYKI-53655 and NBQX (10-20 µM). Blue lines denote the occurrence of light stimulation. (E) Population data showing the absolute amplitude of EPSCs as a function of light stimulus number. (F) Representative current traces of an LOC neuron in response to single light stimulations at the V_H_ from −93 to +47 mV, in steps of 10 mV. Thickened traces represent V_H_ at −93, +7 and +47 mV. (G) Population data showing the I-V relation of EPSC in control and with bath application of MK-801 (10 µM). Inset shows the overlaid, normalized EPSCs at +37 mV under these two conditions. N in (E) and (G), the number of neurons. Error bars in (E) and (G), SEM.

### LOC neurons receive glycinergic inhibitory inputs from MNTB

As the MNTB is a major source of inhibition in the auditory brainstem, a stimulating electrode was placed over MNTB and voltage-clamp experiments were conducted on LOC neurons of the ipsilateral LSO (Figure 7A). At a V_H_ of −30 mV, we observed inhibitory postsynaptic currents (IPSCs) whose amplitude gradually increased when we raised the stimulation intensity from 10 V to 80 V (Figure 7B and 7C). Given the increments in amplitude, we estimate that LOC neurons receive at least 5 inhibitory inputs on average. The IPSC decay time constant was 5.84 ± 0.42 ms (n = 10 neurons). Next, we applied a train of electrical stimulation at frequencies varying from 10 to 200 Hz. Representative IPSCs from one LOC neuron are shown in Figure 7D and 7E. The amplitudes of normalized IPSCs were plotted as a function of stimulus number (Figure 7F). At 10 or 20 Hz, IPSCs exhibited short-term depression (Figure 7D and 7F), while at 200 Hz, the amplitude of IPSCs increased due to summation (Figure 7E and 7F). 50 and 100 Hz stimulation elicited an intermediate pattern in which the amplitude increased at first and then decreased (Figure 7F). Strychnine (500 nM), a glycine receptor antagonist, blocked ~87% of the IPSC, while the remaining current was blocked by gabazine (10 µM), a GABA receptor antagonist (Figure 7G and 7H). Because MNTB neurons are glycinergic, this small GABA current could originate from other inhibitory sources such as superior paraolivary nucleus (SPN, Figure 7A, ref 34). Finally, we varied V_H_ from −50 mV to −70 mV to characterize the I-V relations of IPSC (Figure 7I and 7J). The slope (i.e, inhibitory conductance) of this curve was 1.563 nS, and x-intercept (reversal potential) was −81.5 mV. Thus, LOC neurons receive glycinergic inhibitory inputs from MNTB and possibly nearby nuclei activated by the stimulus.

**Figure 7.**
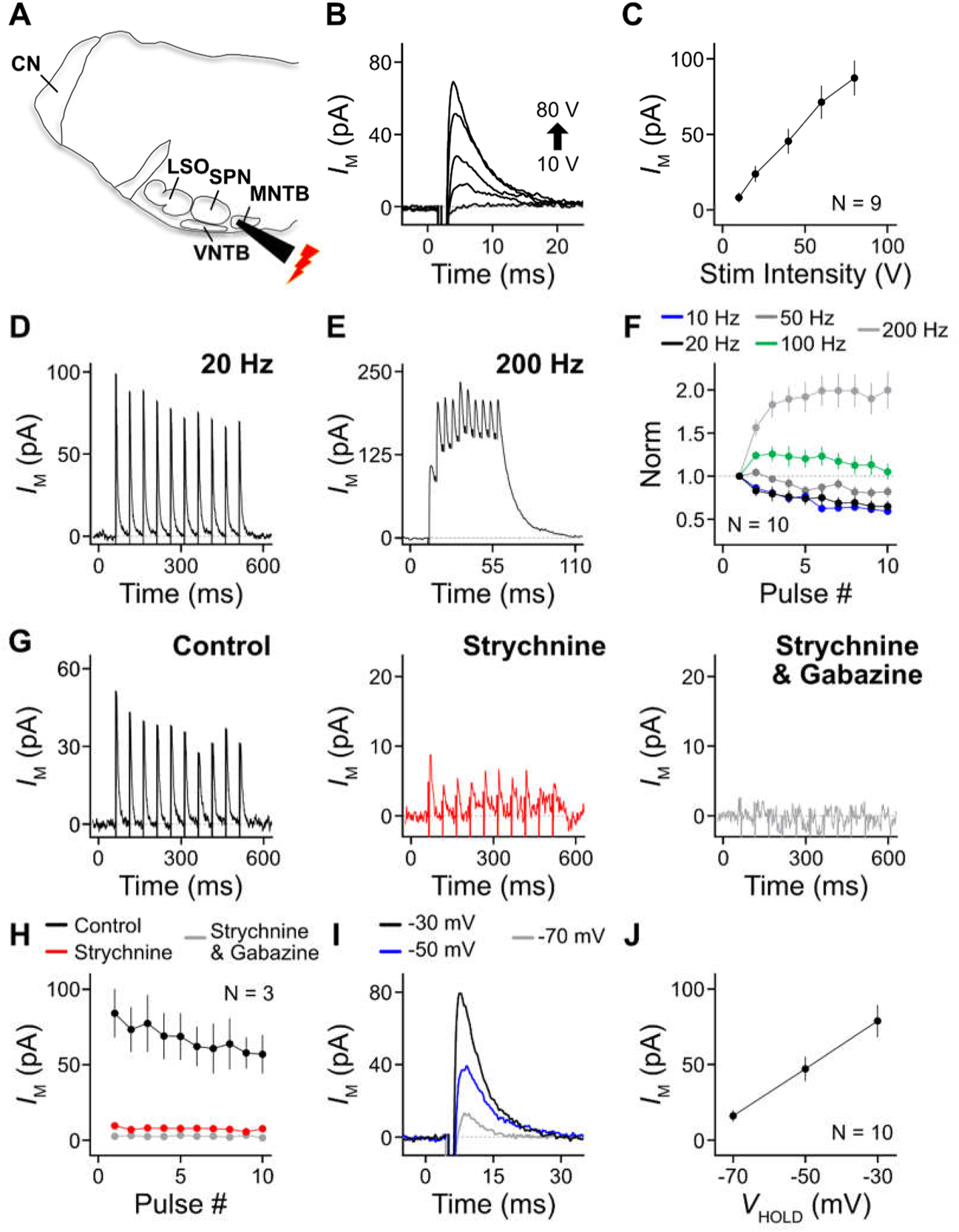
LOC neurons receive glycinergic, inhibitory inputs from the medial nucleus of trapezoid body (MNTB) (A) Schematic figure showing the placement of stimulating electrode over MNTB. (B) Representative current traces of an LOC neuron in response to electrical stimulation with intensity of 10, 20, 40, 60 and 80 V. V_H_ is −30 mV unless otherwise mentioned. Stimulating artifacts were blanked in all representative traces for graphing purposes. (C) Population data showing the amplitude of IPSCs as a function of stimulating intensity. (D-E) Representative current traces of an LOC neuron in response to a train of electrical stimulation at 20 (D) and 200 Hz (E). (F) Population data showing the normalized amplitude of IPSCs (to the amplitude of first IPSC) as a function of electrical stimulus number. (G) Representative current traces of an LOC neuron in response to electrical stimulation of 20 Hz in control, with strychnine (500 nM), and with strychnine and gabazine (10 µM). (H) Population data showing the amplitude of IPSCs as a function of electrical stimulus number, under three different conditions. (I) Representative current traces of an LOC neuron in response to electrical stimulation at V_H_ of −30, −50 and −70 mV. (J) Population data showing the I-V relation of IPSC. N in (C), (F), (H) and (J), the number of neurons. Error bars in (C), (F), (H) and (J), SEM.

### Conductance clamp reveals interaction between intrinsic oscillations and synaptic inputs

After identifying the sources and properties of excitatory and inhibitory inputs to LOC neurons, we explored how they interact with the intrinsic firing mechanism of these neurons described above. Conductance clamp was used to deliver inhibitory (IPSGs, Figure 8A, inset, lower panel) and excitatory (EPSGs, Figure 8B, inset, lower panel) conductances based on the measured amplitudes, decay time constants, and profiles of synaptic depression. Each conductance-clamp train consisted of 20 IPSGs or EPSGs delivered at 20 Hz, a frequency similar to spontaneous activity recorded in CN and MNTB *in vivo* (35, 36). Figure 8A shows an example in which a train of IPSGs was injected every 6 s, with the occurrence of IPSG trains marked by blue triangles. The gray area is enlarged in the inset to show that each IPSG pulse gradually hyperpolarized the membrane voltage and hence stopped the burst (Figure 8A, inset, upper panel). In general, the IPSG train was able to terminate the burst in 81% of trials (total number of trials = 137 from 8 neurons). To further quantify this inhibition, two variables were calculated: *L*_Burst_, representing the length of burst firing, and *T*_IPSG_, representing the timing of the IPSG relative to the onset of the burst (Figure 8A). A histogram of *T*_IPSG_ is shown in Figure 8C, and highlights cases when the burst was successfully inhibited (black) and when it was not (red). All inhibition failures occurred when the IPSGs occurred soon after the onset of the burst (i.e., *T*_IPSG_ < 2 s), indicating that once a burst began, it could not be stopped by these inhibitory stimuli during a critical period. This period in which the burst is refractory to inhibition could be a result of a heightened Ca^2+^ current at the beginning of each burst. However, after this 2-second refractory period, bursts were readily halted by IPSGs; indeed, a plot of *L*_Burst_ as a function of *T*_IPSG_ is strikingly linear (Figure 8D).

**Figure 8.**
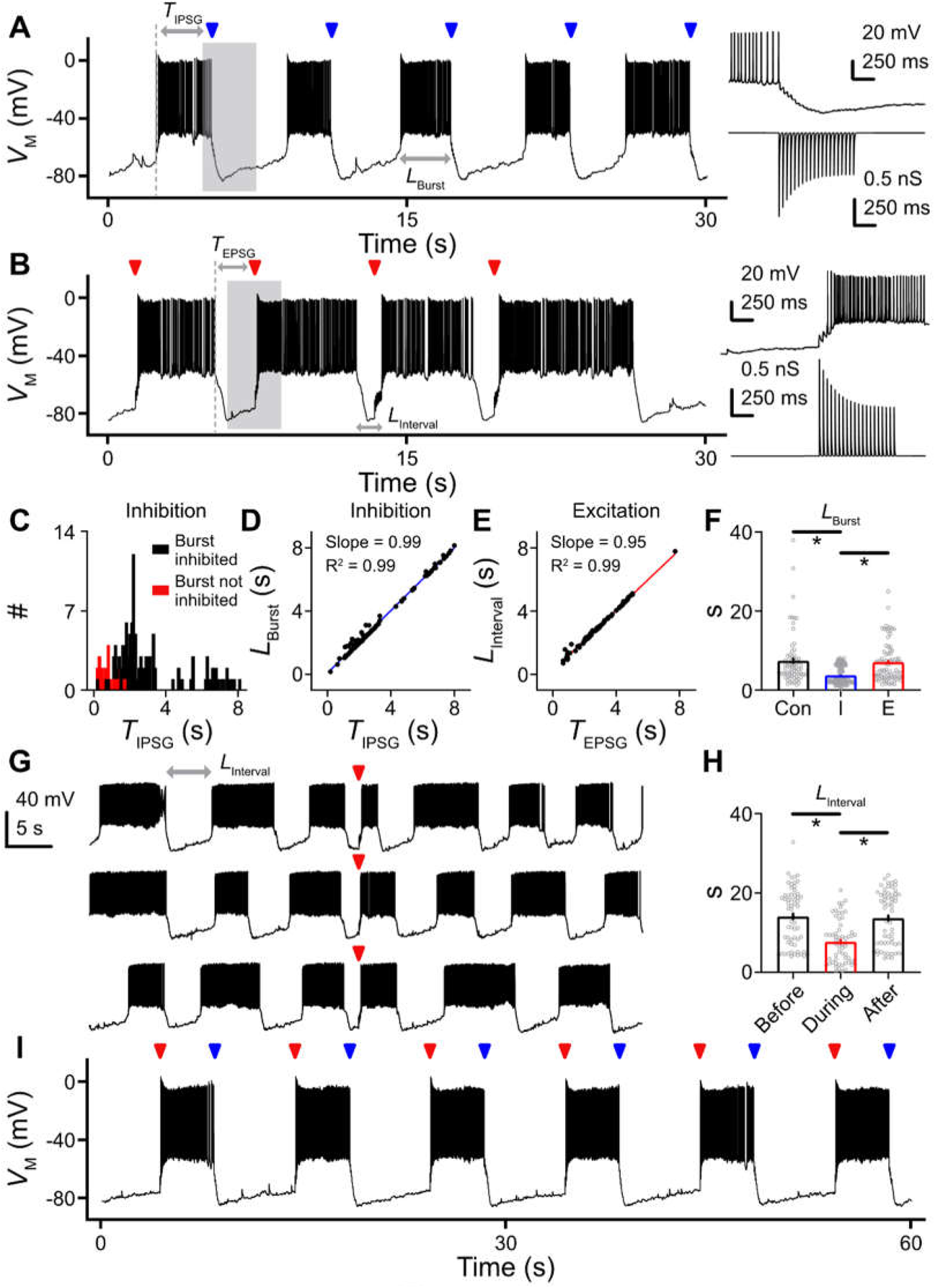
Conductance clamp reveals interaction between intrinsic oscillations and synaptic inputs. (A) Representative voltage traces of an LOC neuron in response to the injection of IPSG train every 6 s. Blue triangles denote the occurrence of IPSG train. Grey area is enlarged in the inset that shows the cessation of a burst (upper panel) induced by the IPSG train (lower panel). Holding current of −5 pA was applied to this LOC neuron. *L*_Burst_, the length of burst. *T*_IPSG_, the timing of IPSG relative to the onset of the burst. (B) Representative voltage traces of the same LOC neuron as in (A) in response to the injection of EPSG train every 6 s. Red triangles denote the occurrence of EPSG train. Grey area is enlarged in the inset that shows the generation of a burst (upper panel) induced by the EPSG train (lower panel). Holding current of −5 pA was applied. *L*_Interval_, the length of inter-burst interval. *T*_EPSG_, the timing of EPSG relative to the onset of the interval. (C) Histogram of *T*_IPSG_ showing two categories, when the burst was successfully inhibited (black) and when it was not (red). (D) Scatter plot showing *L*_Burst_ as a function of *T*_IPSG_. Blue line represents a linear fit to the data points. (E) Scatter plot showing *L*_Interval_ as a function of *T*_EPSG_. Red line represents a linear fit to the data points. (F) Population data showing *L*_Burst_ in control (con), with IPSG (I) or EPSG (E) injection. * = *p* < 0.05 (*post hoc* Bonferroni adjusted *t* test). Each dot represents a trial, data from 8 neurons. (G) Three-minute continuous recording from a representative LOC neuron when the EPSG train was applied every 60 s. Red triangles denote the occurrence of EPSG train. Holding current of −20 pA was applied to this LOC neuron. (H) Population data showing *L*_Interval_ before, during and after the application of EPSG train. * = *p* < 0.05 (*post hoc* Bonferroni adjusted *t* test). Each dot represents a trial, data from 7 neurons. Error bars in (F) and (H), SEM. (I) Representative voltage traces of the same LOC neuron as in (A) and (B) in response to the injection of EPSG (red triangle) paired with IPSG (blue triangle). The interval between EPSG and IPSG was set at 4 s, and the duration of each period of synaptic stimulation was 1 second. Holding current of −15 pA was applied.

We then injected trains of EPSGs to the same set of LOC neurons (Figure 8B, red triangles). EPSGs appearing during the quiet period between bursts summated, and within a few EPSGs triggered a long spike burst in 98% of trials (Figure 8B, inset, upper panel; total number of trials = 82 from the same 8 neurons). Similar to the analysis of IPSGs, we calculated two variables: *L*_Interval_, the length of inter-burst interval, and *T*_EPSG_, the timing of EPSG relative to the onset of the interval. The relationship between *L*_Interval_ and *T*_EPSG_ yielded a slope close to 1, indicating that bursts were reliably triggered by the EPSGs (Figure 8E).

By comparing *L*_Burst_ across three conditions (i.e., control, IPSG injection and EPSG injection), we found that inhibition significantly curtailed the duration of bursts (Figure 8F, *p* = 0.018, Bonferroni *post hoc* test, n = 8 neurons). Because burst length was shortened, the neurons were now able to generate bursts at a higher frequency (from 0.07 Hz to 0.13 Hz, *p* = 0.001, Bonferroni *post hoc* test). By contrast, excitation did not change the length of bursts (Figure 8F, *p* > 0.9999, Bonferroni *post hoc* test). These observations suggest that excitation functions to reset the bursting activity (i.e., determine the onset of the burst), but not to alter its rhythm. We thus applied EPSG trains every 60 s, illustrated by a 3-minute continuous recording in Figure 8G. The *L*_Interval_ before and after the application of an EPSG train were not significantly different (Figure 8H, *p* = 0.620, Bonferroni *post hoc* test, n = 60 trials from 7 neurons), but the *L*_Interval_ at which the EPSGs occurred was curtailed by the evoked bursts (*p* < 0.0001, Bonferroni *post hoc* test). Taken together, synaptic inhibition of LOC neurons increases the pace of bursting activity while synaptic excitation reset the timing of this activity. By combining the distinct synaptic functions, the pattern of bursting activity in LOC neurons could be uniquely tailored by excitation and inhibition (Figure 8I). Here, *L*_Burst_ was 3.99 ± 0.03 s, nearly identical to the 4-s interval between EPSG and IPSG (*p* = 0.643, paired *t* test, n = 64 trials from 5 neurons), indicating that synaptic stimuli switched on and off the spike bursts.

## Discussion

We identified a mechanism for extremely slow electrical signaling in the LOC system. This low rate of patterned activity is uncommon in the auditory brainstem, where speed and precision are traditionally believed to be the currency of signaling (37). Underlying this activity is an L-type Ca^2+^ channel-dependent intrinsic oscillator that gives rise to long trains of Na^+^ channel-dependent APs. This activity is further guided by synaptic inputs. Overall, the interaction between synaptic input and the intrinsically generated excitation leads to a novel signaling mode, in which relatively brief synaptic excitation serves to turn on the spike train, and brief synaptic inhibition then turns it off, after a minimal firing period of about 2 seconds. The source of excitation originates from the distal ends of the central auditory pathway: a bottom-up input pathway from T-stellate cells in the CN converges with a top-down pathway from pyramidal cells of the auditory cortex. GluA2-containing AMPA receptors mediate transmission at both synapses while some cortical inputs also engage kainate receptors. Inhibition to LOC neurons is supplied by glycinergic MNTB neurons. As discussed below, the prolonged period of burst firing in LOC neurons may play an essential role in releasing a diverse cohort of neurotransmitters and neuromodulators (particularly peptides) in the cochlea, while the convergent inputs to LOC neurons may enable them to adjust their output in response to changing sensory environments.

### Oscillations are age- and Ca^2+^-dependent

The activity of LOC neurons has not been described *in vivo* as the LOC axons are small and unmyelinated, and thus their signals are easily obscured by those of myelinated afferent and efferent (MOC) fibers (38–40). A previous study of identified LOC neurons recorded *in vitro* reported no spontaneous activity (15). However, that study only looked at rats at ages P5-11, prior to hearing onset. Our examination of P9-11 mice confirmed that the burst pattern we report in older mice is absent before hearing onset. Other features of these young neurons were similar in the two species, including an apparent lack of HCN current (i.e., voltage sag upon hyperpolarization), and the presence of a delay in the onset of spiking characteristic of A-type K^+^ current. These shared features of LOC neurons were also recognized in other studies in which no biomarker was used to distinguish LOC neurons from their neighbors (16, 17, 41). Therefore, it seems likely that the intrinsic, spontaneous electrical activity of LOC neurons arises from an age-, and possibly experience-, dependent mechanism. In line with this, we found that spontaneous activity starts to emerge at P9-11 in a small population of LOC neurons (~44%), but manifests as tonic firing instead of a burst firing pattern. Interestingly, Ca^2+^ channels in prehearing neurons require an activation voltage more positive than the threshold of AP (15). With a high activation voltage, it is unlikely that these Ca^2+^ channels would constitute an intrinsic oscillator. In contrast, burst firing pattern was readily found in >90% of LOC neurons in juvenile and young adult mice, suggesting that the development of Ca^2+^ channels or other components of oscillatory behavior may appear between P11-17. However, while the prehearing LOC neurons lack the mature intrinsic oscillator that drives infra-slow burst activity, there may still be burst activity in prehearing LOC neurons *in vivo*, due to the spontaneous activity that originates in the immature cochlea and radiates throughout the immature central auditory system (42, 43). Indeed, developmental loss of this source of activity might drive the emergence of an intrinsic oscillator in LOC neurons.

### A complex neural circuit interconnects bilateral control of LOC neurons

Previous studies found that both bushy and T-stellate cells in the CN project to the LSO (27, 44). In contrast to the well-characterized connections between bushy cells and LSO principal cells, the target of T-stellate cells in the LSO has remained elusive. We show here that T-stellate cells form the ascending input pathway to the ipsilateral LOC neurons. While this connection from the ipsilateral CN completes the canonical LOC reflex pathway, from cochlea to CN to LOC and back to cochlea, our results suggest a much more complex circuitry and function. First, MNTB neurons, which receive inputs from bushy cells in the contralateral CN, project to LOC neurons, indicating that activity from the contralateral ear controls LOC neurons through synaptic inhibition and control of the burst cycle. Second, the ipsilateral auditory cortex sends descending excitatory inputs to LOC neurons. This input contains a representation of activity from the contralateral ear. Thus, while the canonical reflex path of LOC activity is monaural (1), in fact the spike activity in LOC neurons is under binaural control. Cortical signaling could also relay non-auditory information from other modalities. Interestingly, a stress-induced tinnitus model suggests a key role of LOC neurons releasing opioid peptides in the inner ear (12); it is possible that cortical input participates in this signaling.

Some descending inputs utilized kainate receptors in addition to AMPA receptors, and consistent with previous reports, featured slow-gating kinetics that lead to summating plateau responses (45–48). Although the current was small, its slow time course, in combination with the high input resistance of the neuron, likely leads to a substantial depolarization. Kainate current-mediated connections was higher with viral injection into IC than that into the auditory cortex, and this difference could result from higher infection by retrograde virus (injected into the IC) versus anterograde virus (injected into the auditory cortex). Indeed, the AMPA current obtained with IC injection was also larger than that with cortex injection (compare Figure 5D and 6E, EPSCs at 20 Hz, *p* = 0.009, n = 15 in 5D and 7 neurons in 6E, Mann-Whitney test). Therefore, we expect a higher proportion of kainate-receptor containing synapses *in vivo*.

### Functional significance of spike burst activity in LOC neurons

For decades, the LOC system, along with its peptidergic modulation, has been thought to play a fundamental role in modulating normal hearing function. LOC neurons express a plethora of neurotransmitters and neuromodulators, varying from low molecular-weight neurotransmitter such as acetylcholine, GABA and dopamine, to high molecular-weight neuropeptides including CGRP, urocortin, dynorphin and enkephalin (1, 10). It was proposed that dynorphin and CGRP have an excitatory effect, enhancing the neural activity of the cochlea and thus improving the signal to noise ratio, while enkephalin is inhibitory (49–52). In mice lacking αCGRP, sound-evoked activity was reduced in the auditory nerve (53). In addition, a tinnitus model suggested that dynorphins released by LOC neurons could increase the sensitivity of NMDA receptors in the auditory nerve terminals, amplifying signals from inner hair cells and enhancing auditory nerve activity. Chronic exposure to dynorphins (e.g., stress induced) could result in deleterious effects including tinnitus (12). Urocortin and its receptors control cochlear sensitivity and protect the ear from acoustic trauma. Mice lacking urocortin displayed elevated thresholds of both auditory brainstem response and otoacoustic emission (13). Knocking out urocortin receptors increased the susceptibility to noise-induced hearing loss (14).

The electrical activity in the LOC system appears optimized for these important functions of peptide signaling in the cochlea. The frequency of brain oscillations varies widely, from γ oscillations with a frequency band between 30-80 Hz (54), to infra-slow oscillations of <0.1 Hz found in thalamic relay nuclei (55, 56) and in the LOC system. These slowest of oscillations are characterized by long periods of firing and silence. Such long stretches of firing may be necessary for the releasing of peptides in the cochlea, as has been observed in hypothalamus, neurohypophysis and hippocampus (for review see 57, 58, 59). Those studies suggest that Ca^2+^-dependent release of a peptide-containing vesicle is a rare event triggered only after prolonged membrane depolarization or hundreds of presynaptic spikes (57, 60). Release of oxytocin, vasopressin and brain-derived neurotrophic factor was more effective when spike bursts were followed by intervals of silence (61–64), which may accommodate vesicle replenishment. These conditions for peptide release are met by the intrinsic firing activity of LOC neurons.

Synaptic input to LOC neurons may affect peptide release in multiple ways. The triggering and cessation of bursts by synaptic input will impact the likelihood of peptide release and the time for recovery of vesicles. The projection from CN to LSO is tonotopically organized (23), suggesting that sound onset will synchronize spike bursts in nearby LOC neurons; this could result in a bolus of peptide release in the cochlea from a population of efferent fibers. Moreover, recent observations show that LOC neurons respond dynamically to auditory stimuli by regulating the expression of dopamine and possibly other transmitters (65). Therefore, synaptic input to LOC neurons could control both the timing of exocytosis and vesicle content.

## Materials and Methods

**See Supplemental Information**

## Acknowledgments

This work was supported by NIH grant DC004450 to L.O.T. We thank Doug Zeppenfeld for assistance with imaging, and Dr. Gabriel Romero for comments on the manuscript.

## Supplementary Information

### Supplementary Information Text

#### Materials and Methods

##### RESOURCE AVAILABILITY

###### Lead contact

Further information and resources should be directed to and will be fulfilled by the lead contact, Laurence Trussell.

###### Materials availability

This study did not generate new unique reagents.

###### Data and code availability

Electrophysiology and microscopy data reported in this paper will be shared by the lead contact upon request.

##### EXPERIMENTAL MODEL AND SUBJECT DETAILS

###### Mice

Mice were maintained in the animal facility managed by the Department of Comparative Medicine at Oregon Health and Science University. All procedures were approved by the Oregon Health and Science University’s Institutional Animal Care and Use Committee and met the recommendations of the Society of Neuroscience. Transgenic mice of both sexes expressing Cre recombinase under the endogenous choline acetyltransferase promoter (ChAT-IRES-Cre; Jackson Labs 031661) were used. Except for calcium imaging experiments, these mice were crossed with a tdTomato reporter line (Ai9(RCL-tdT); Jackson Labs 007909) to generate mice that express tdTomato in cholinergic neurons (referred to as ChAT-Cre/tdTomato).

##### METHOD DETAILS

###### Acute brainstem slice preparation

Juvenile and young adult mice at postnatal days (P) 17-42 were deeply anesthetized with isoflurane and decapitated. To characterize the age-dependent difference in biophysical properties of LOC neurons, prehearing mice at P9-11 were also used. The brain was dissected in ice-cold (0°C) low-sodium artificial cerebral spinal fluid (aCSF) containing (in mM) 260 glucose, 25 NaHCO_3_, 2.5 KCl, 1.25 NaH_2_PO_4_, 1.2 CaCl_2_, 1 MgCl_2_, 0.4 ascorbic acid, 3 myo-inositol and 2 sodium pyruvate, continuously bubbled with a mixture of 95% O_2_/5% CO_2_. The brain was blocked coronally, affixed to the stage of a vibratome slicing chamber (VT1200S, Leica) and submerged in low-sodium aCSF. Coronal brain slices containing the superior olivary complex (SOC), cochlear nucleus (CN) and inferior colliculus (IC) were cut at 300 µm and transferred to normal aCSF at 34°C containing (in mM) 125 NaCl, 25 NaHCO_3_, 2.5 KCl, 1.25 NaH_2_PO_4_, 1.2 CaCl_2_, 1 MgCl_2_, 10 glucose, 0.4 ascorbic acid, 3 myo-inositol and 2 sodium pyruvate (pH 7.4, osmolarity 300-310 mOsm/l). Normal aCSF was continuously bubbled with a mixture of 95% O_2_/5% CO_2_. When sectioning was completed, slices were incubated for an additional 30 min at 34°C, followed by storage at room temperature, ~22°C.

###### Electrophysiology

Upon electrophysiological experiments, slices were transferred to a recording chamber mounted on a Zeiss Axioskop 2 FS plus microscope. The microscope was equipped with a CCD camera, 10× and 40× water-immersion objectives. The recording chamber was perfused with normal aCSF at 3 ml/min and maintained at 31–33°C with an in-line heater (TC-324B; Warner Instrument Corp). Neurons in each slice were viewed using full-field fluorescence with a white-light LED attached to the epifluorescence port of the microscope that was passed through a tdTomato filter set. LOC neurons expressing tdTomato were identified in the lateral superior olive (LSO) that displayed a typical “S” shape on the coronal brainstem slice (Figure 1B).

Borosilicate glass capillaries (OD 1.5 mm; World Precision Instruments) were pulled on a PP-830 Narishige vertical puller to a tip resistance of 2-6 MΩ for cell-attached and whole-cell voltage-clamp experiments, and of 6-10 MΩ for whole-cell current- and conductance-clamp experiments. For cell-attached experiments, recording pipette was filled with normal aCSF. All of whole-cell current- and conductance-clamp experiments were performed with an internal pipette solution containing (in mM) 113 K-gluconate, 2.75 MgCl_2_, 1.75 MgSO_4_, 9 HEPES, 0.1 ethylene glycol tetraacetic acid (EGTA), 14 tris-phosphocreatine, 0.3 tris-GTP, 4 Na_2_-ATP, pH adjusted to 7.2 with KOH, and osmolality adjusted to 290 mOsm with sucrose. In some experiments, 1% biocytin was added to the internal solution for *post hoc* identification of LOC neurons. Whole-cell voltage-clamp experiments were conducted using this K^+^-based internal solution or a Cs^+^-based internal solution containing (in mM) 103 CsCl, 10 tetraethylammonium chloride (TEA-Cl), 3.5 N-ethyllidocaine chloride (QX-314-Cl), 2.75 MgCl_2_, 1.74 MgSO_4_, 9 HEPES, 0.1 EGTA, 14 tris-phosphocreatine, 0.3 tris-GTP, 4 Na_2_-ATP, with pH adjusted to 7.2 with CsOH, and osmolality adjusted to 290 mOsm with sucrose. All of the experiments that aimed to characterize the current-voltage relations of excitatory synaptic current were conducted with this Cs^+^-based internal solution. In a subset of experiments (n = 4 neurons), 0.1 mM spermine was added to the internal solution to compensate the dialysis of endogenous polyamines. Recording pipettes were wrapped with Parafilm M (Bemis) to reduce pipette capacitance. Junction potential was experimentally measured as −10 mV and −3 mV for K^+^- and Cs^+^-based internal solution, respectively. All of our data were reported with the correction of junction potentials.

Electrophysiological experiments were performed using an Axon Multiclamp 700B amplifier (Molecular Devices), except for conductance-clamp experiments, which were performed using a dPatch amplifier (Sutter Instrument). Raw data were low-pass filtered at 10 kHz and digitized at 100 kHz using a Digidata 1440A (Molecular Devices). Recording pipettes were visually guided to the LSO region and membrane patches were ruptured after a GΩ seal was attained. In current-clamp mode, LOC neurons were initially silent upon break-in, but all of them became spontaneously active within 5 minutes after whole-cell configuration. In some cases, up to −30 pA bias current was applied if a neuron became excessively depolarized. Optogenetic experiments were performed in voltage-clamp mode with the presence of strychnine (500 nM) and gabazine (10 µM). Series resistance compensation was set to 70% correction and prediction. LOC neurons were held at −70 mV (K^+^-based internal) or −63 mV (Cs^+^-based internal) while channelrhodopsin (ChR2) expressed at presynaptic terminals was activated using 2-ms light flashes through a GFP filter set from a 470 nm LED attached to the epifluorescence port of the microscope. A train of light simulation at 5-20 Hz was applied to LOC neurons every 30 s, and each frequency contained at least 10 trials. Excitatory postsynaptic currents (EPSCs) were averaged over these trials for each simulation frequency. To profile the current-voltage relations of EPSCs, single light stimulations were employed when an LOC neuron was held at membrane potentials from −93 mV to +47 mV in steps of 10 mV. The amplitude of EPSCs was measured and plotted as a function of membrane potential. In order to increase the release probability of the presynaptic terminal, 4-AP (50 µM) was applied in the bath in the experiments with viral injections into IC or auditory cortex. In a subset of experiments with viral injection into CN (n = 7 neurons), 4-AP was also used to try to uncover a kainate receptor component by boosting release probability. GYKI-53655 (50 µM), NBQX (10-20 µM) and MK-801 (10 µM) were applied to block AMPA, kainate and NMDA receptors, respectively. To study short-term plasticity of the synapses between CN and LOC, a stimulating electrode was placed in between CN and LSO to excite CN fibers. We also exploited electrical stimulation to investigate the inhibitory inputs to LOC neurons, by placing the stimulating electrode over the medial nucleus of trapezoid body (MNTB). With the presence of NBQX (10 µM), LOC neurons were held at −30 mV (K^+^-based internal) while stimulating intensity was raised gradually from 10 V to 80 V. At 10 V, the amount of inhibitory postsynaptic current (IPSC) was minimal. Next, the stimulating intensity was set at 60-80 V while a train of electrical pulses from 10 Hz to 200 Hz was applied every 10 s; each frequency contained at least 10 trials. IPSCs were averaged over these trials for each stimulation frequency. To profile the current-voltage relations of IPSCs, holding voltage was lowered to −50 mV and −70 mV. Strychnine (500 nM) and gabazine (10 µM) were used to block glycine and GABA receptors, respectively.

In a subset of experiments (n = 5 neurons), gramicidin-perforated patch-clamp recordings were employed to characterize the spontaneous burst firing of LOC neurons instead of whole-cell recordings (1). Gramicidin was diluted in the aforementioned K^+^-based internal solution at 50 µg/mL. Recording pipettes had a resistance of 6-10 MΩ. The tip of recording pipette was first filled with gramicidin-free internal solution and then back-filled by gramicidin-containing solution. After forming a GΩ seal, the degree of perforation was monitored in current-clamp mode. Because LOC neurons were spontaneously active, the progression of perforation could be reflected by the growth of spike amplitude. We started recording ~45 minutes after, as the series resistance dropped to ~30 MΩ or less. Bursting activity of LOC neurons recorded from perforated patch resembles that from whole-cell current-clamp experiments. Therefore, these two sets of data were pooled.

###### Conductance-clamp protocols

Conductance-clamp experiments were performed using a dPatch amplifier (Sutter Instruments). Raw data were low-pass filtered at 10 kHz and digitized at 50 kHz. Excitatory and inhibitory conductance waveforms (EPSG and IPSG, respectively) were created in Igor Pro 8 based on our physiological data. The unitary PSG waveform was constructed as

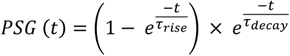

where τ_rise_ = 0.7 ms for both EPSG and IPSG; τ_decay_ = 4.27 ms for EPSG and 5.84 ms for IPSG. A train of PSGs at 20 Hz for 20 pulses (Figure 8A inset and 8B inset) was constructed by scaling the initial PSG to values obtained for low-frequency responses (1.786 nS for EPSG and 1.563 nS for IPSG). Reversal potential of the excitatory and inhibitory conductances were 0 mV and −80 mV, respectively. Depression was approximated by scaling down subsequent PSGs in the train according to a “time constant” of four stimuli (200 ms for a 20 Hz train) to an equilibrium level of 0.5 times the initial PSG value.

###### Calcium imaging

Acute brainstem slices from mice with viral injection into the LSO were harvested as described above 1-2 weeks after the surgery. A two-photon imaging system (Prairie Technologies) powered by a Ti:sapphire pulsed laser (Chameleon Ultra II) was used for calcium imaging. The laser was tuned to 1000 nm. Epi- and transfluorescence signals were captured through a 20×, 0.5 NA objective and a 1.4 NA oil immersion condenser (Olympus). Fluorescent signals were split into red and green channels using dichroic mirrors and band-pass filters (epi: 575 DCXR, HQ525/70, HQ607/45; trans: T560LPXR, ET510/80, ET620/60; Chroma). Red fluorescence (mRuby) was captured with R9110 photomultiplier tubes (PMTs, Hamamatsu). Green fluorescence (GCaMP6f) was captured with H8224 PMTs. Data were collected in frame-scan mode (0.5754s/frame). Each LOC neuron represents a single region of interest (ROI). For individual ROIs, the intensity of fluorescence was measured in Fiji and F_G_/F_R_ was calculated as the intensity of green signals divided by the intensity of red signals. For comparison purposes across multiple LOC neurons, this ratio was further scaled to a range between 0% and 100% (termed as scaled F_G_/F_R_) based on the following formula and plotted as a function of time (see Figure 2):

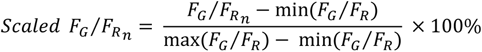

In order to define whether an LOC neuron was spontaneously active and its oscillating frequency, scaled F_G_/F_R_ data were further processed in Igor Pro 8. First, the histogram of scaled F_G_/F_R_ was plotted for each neuron. If Ca^2+^ signals of an LOC neuron oscillated with alternating peaks and troughs, the histograms would show bimodal distribution (Figure S2A and S2B). The lower peak represents the troughs, i.e., the floor of the baseline. A Gaussian fit was applied to the lower peak in order to derive the mean and standard deviation (SD) of the baseline (Figure S2B). The threshold was defined as the mean plus 4 SD (Figure S2A, red dash line). LOC neurons were categorized as spontaneously active if their signals exceeded this threshold. Second, the power spectrum of scaled F_G_/F_R_ was plotted and the oscillation frequency was defined as the frequency at which the correlation coefficient exhibited an initial peak (Figure S2C, arrowhead).

###### Immunohistochemistry and confocal imaging

Mice were deeply anesthetized with isoflurane and then transcardially perfused with 0.1 M phosphate buffered saline (PBS) followed by 4% paraformaldehyde in 0.1 M PBS using a peristaltic pump. Brains were surgically extracted and incubated in 4% paraformaldehyde in 0.1 M PBS overnight at 4°C. To visualize the LOC neurons filled with biocytin during whole-cell recording, 300 µm acute brainstem slices were fixed with 4% paraformaldehyde in 0.1 M PBS overnight at 4°C. On the next day, brains or brainstem slices were rinsed three times in 0.1 M PBS, 10 minutes per rinse. Then brains were embedded in 4% agar and affixed to the stage of a vibratome slicing chamber and then sectioned in the coronal plane at 50 µm. Each section was collected in 0.1 M PBS. For immunostaining, free-floating sections were permeabilized and blocked in 2% bovine serum albumin, 2% fish gelatin, and 0.1% Triton X-100 in 0.1 M PBS for 2 hours at room temperature on a 2-D rocker. Sections were then incubated in primary antibodies with varying concentration for 2 days at 4°C on a 2-D rocker, followed by three rinses in 0.1 M PBS, 10 minutes per rinse. Then the sections were incubated in secondary antibodies and streptavidin-conjugated fluorophores overnight at 4°C on a 2-D rocker. See “Key Resources” table for a full list of antibodies and corresponding concentration we used. Sections were rinsed three times, 10 minutes per rinse on the next day, before being mounted on microscope slides and cover slipped with Fluoromount-G (Southern Biotech) mounting medium, and then sealed with clear nail polish. Some brain sections with high fluorophore expression were not enhanced with antibody labeling to reduce background (e.g., tdTomato in Figure 1C). Images were acquired on a Zeiss LSM780 or LSM900 confocal microscope system. Images were processed for contrast and brightness in Fiji. Neurons filled with 1% biocytin were reconstructed in Neurolucida (MBF Bioscience) by manually tracing the dendritic processes.

###### Stereotactic injections of virus

Glass capillaries (WireTrol II; Drummond Scientific) were pulled on a P-97 Flaming/Brown micropipette puller (Sutter Instrument) and beveled to 45-degree angle with a tip diameter of 40– 45 mm using a diamond lapping disc (0.5 mm grit, 3M). Mice (P21-23) were anesthetized with 4% isoflurane and secured in a small stereotaxic frame (David Kopf). Isoflurane was continuously applied at 1.5% to keep the mice anesthetized during the surgery. The rostral-caudal axis of the head was leveled by adjusting the bregma and lambda into the same focal plane. The lateral-medial axis was leveled by adjusting the spots 2 mm lateral from lambda on the left and right skull into the same focal plane. A unilateral craniotomy was conducted using a dental drill (Foredom K.1070). The ready-made glass pipette was secured in a hydraulic injector (MO-10, Narishige) attached to a triple-axis motorized manipulator (IVM Triple, Scientifica). The manipulator was used to navigate the pipette to the injection site and to advance it into and out of the brain at a speed of ~10 µm/s, while the injector was used to deliver the virus at varying volume dependent on the target structure. There was a 5-minute wait time at the injection site before and after the viral injection. For optogenetics, ChR2-expressing virus was injected into CN, IC or auditory cortex of ChAT-Cre/tdTomato mice. The coordinates for CN injection are (in mm from lambda): 0.17 caudal, −2.43 lateral and 4 deep. 50 nl anterograde-transported virus we used to infect CN is AAV1-CAG-ChR2-Venus-WPRE-SV40 (2). Three injections, 100 nl per injection of the same anterograde AAV were made in the auditory cortex with the coordinates (in mm from lambda) 2.54, 2.04, 1.54 rostral, −3.75 lateral and 1 deep. The coordinates for IC injection are (in mm from lambda): 0.3 caudal, −1.1 lateral and 1 deep. 300 nl retrograde-transported virus we used to infect IC is AAVrg-Syn-ChR2-GFP (3). For calcium imaging, a Cre-dependent GCaMP6f-expressing virus was injected into the LSO of ChAT-Cre mice. The coordinates are (in mm from lambda) 0.5 rostral, −1.2 lateral and 5.3 deep. 100 nl anterograde-transported virus we used to infect LSO is AAV1-CAG-Flex-mRuby2-GSG-P2A-GCaMP6f-WPRE-pA (4). Experiments were conducted 1-2 weeks after the surgery.

##### QUANTIFICATION AND STATISTICAL ANALYSIS

Electrophysiological data were collected using Clampex acquisition and Clampfit analysis software (version 10.9, Molecular Devices). Calcium imaging data were analyzed using Fiji and Igor Pro 8 (Wavemetrics). Statistical analyses and graphing protocols were performed using GraphPad Prism 9. Student *t* tests or analysis of variance (ANOVA) with *post hoc* Bonferroni adjusted *t* tests were used to determine significance, unless otherwise mentioned. The standard for significant differences was defined as *p* < 0.05. Numeric values are reported as mean ± standard error of the mean (SEM). The layout of figures was created with Affinity Designer (Serif).

---

**Figure S1.**
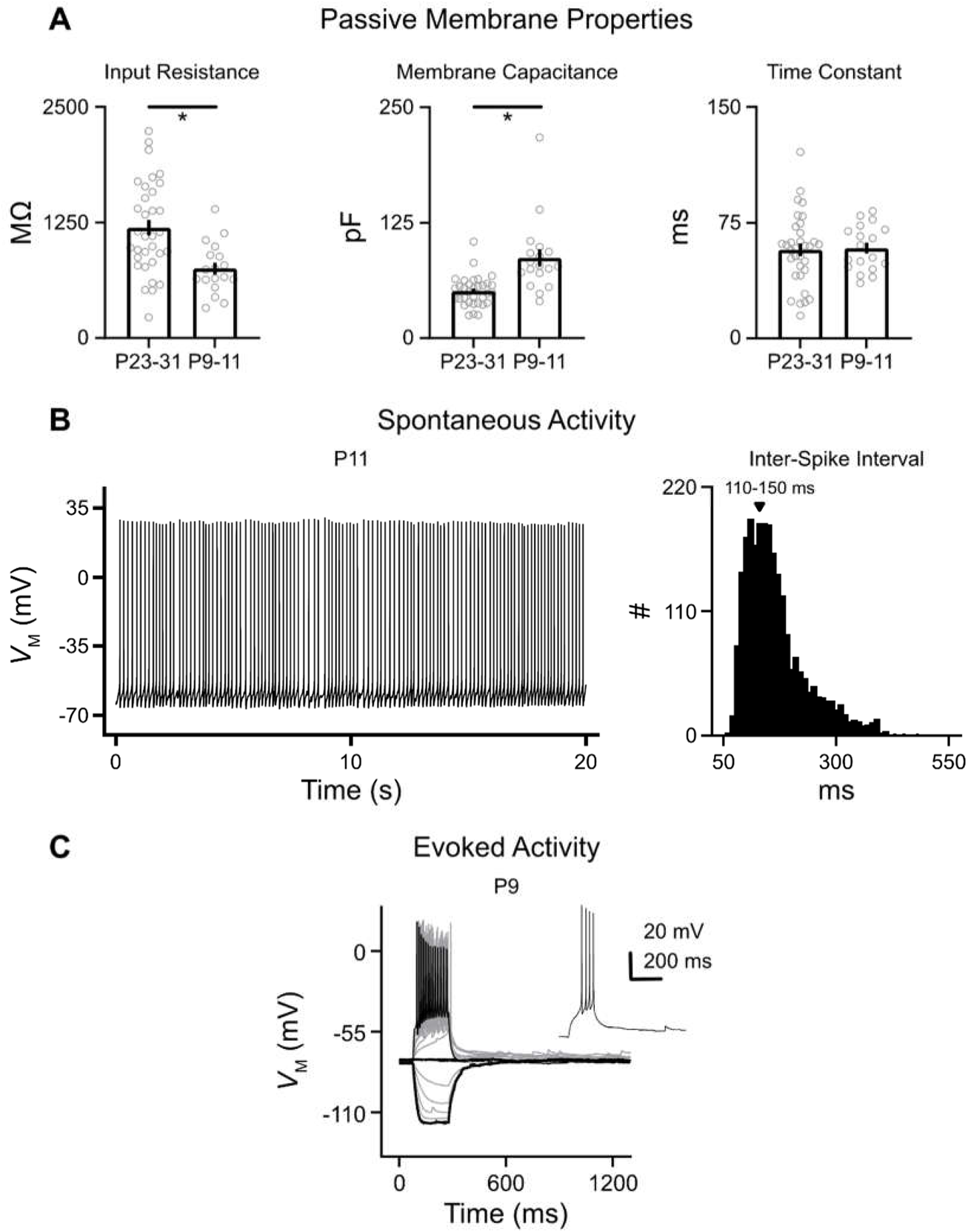
Intrinsic properties of LOC neurons in prehearing mice. (A) Population data showing the passive membrane properties in two age groups. * = *p* < 0.05 (Student *t* test). (B) Left panel: representative spontaneous activity of an LOC neuron at P11, observed in 44% of neurons. Right panel: the histogram of inter-spike interval at P9-11. (C) Representative voltage traces of an LOC neuron at P9 in response to current injections from −100 to 200 pA in steps of 20 pA. Thickened traces represent voltage response to 200, 0 and −100 pA current injections. Inset shows the voltage response to 40 pA current injection.

**Figure S2.**
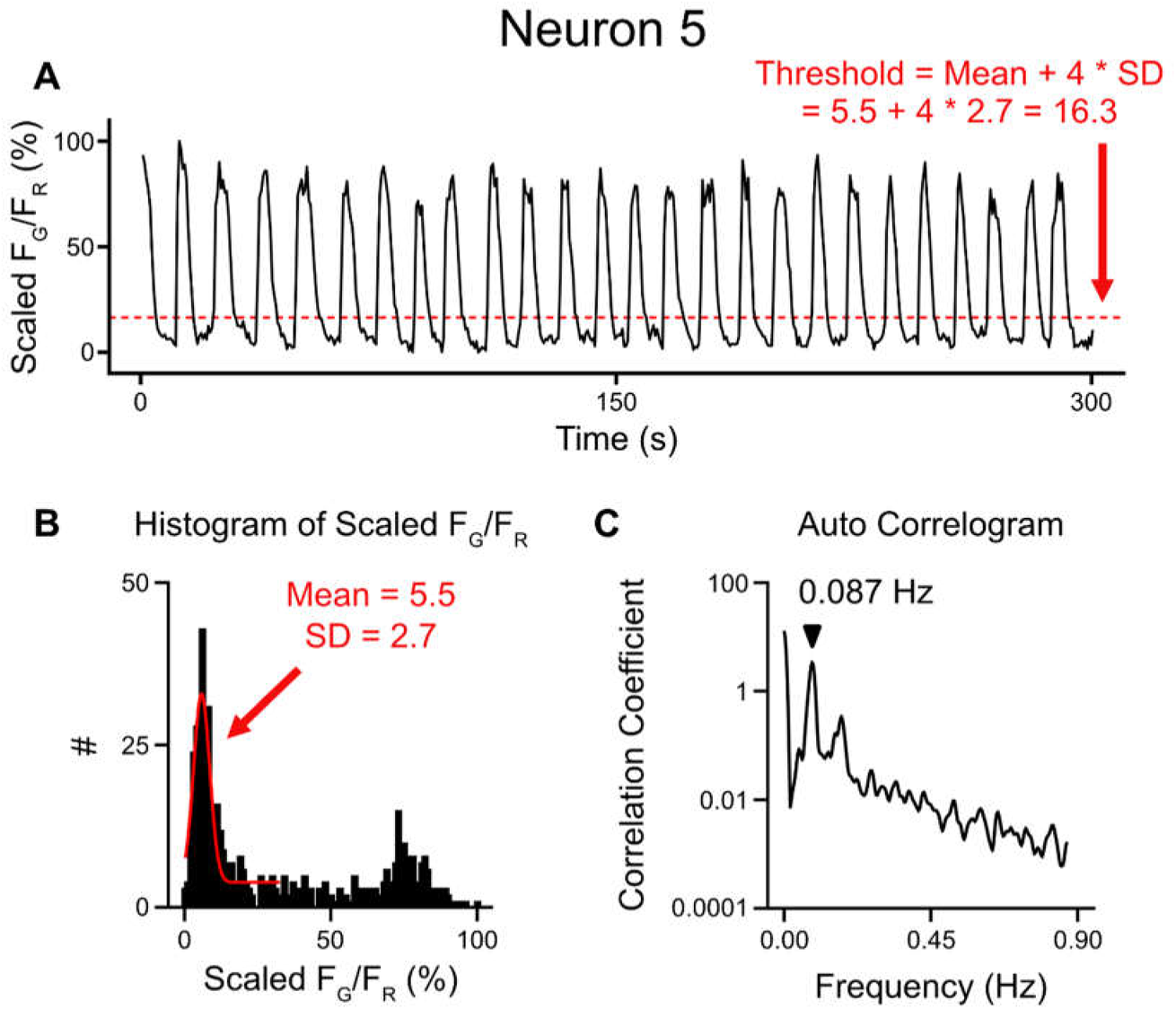
Example of data analysis regarding calcium imaging. (A) Scaled F_G_/F_R_ of Neuron 5 shown in Figure 2C is replotted with red dash line representing the threshold that defines whether a neuron is spontaneously active or not. Mean and standard deviation (SD) is acquired based on the calculation shown in (B). (B) The histogram of scaled F_G_/F_R_ of Neuron 5. A Gaussian fit (red curve) to the lower peak that represents the trough of signals renders the mean of 5.5% and SD of 2.7%. (C) The results of auto-correlation of scaled F_G_/F_R_ for the same neuron. Arrowheads point to the peak at 0.087 Hz (i.e., oscillation frequency).

**Table S1.** KEY RESOURCES TABLE.

| **Reagent type**<br>**(species) or**<br>**resource** | **Designation** | **Source or**<br>**reference** | **Identifier** | **Additional**<br>**information** |
| --- | --- | --- | --- | --- |
| **Strain, strain**<br>**background**<br>**(M. musculus)** | ChAT-IRES-Cre | Jackson<br>Laboratory<br>PMID: 21284986 | RRID: IMSR_JAX:<br>031661 |  |
| **Strain, strain**<br>**background**<br>**(M. musculus)** | Ai9(RCL-tdT) | Jackson<br>Laboratory<br>PMID: 22446880 | RRID: IMSR_JAX:<br>007909 |  |
| **Strain, strain**<br>**background**<br>**(M. musculus)** | C57BL/6J | Jackson<br>Laboratory | RRID: IMSR_JAX:<br>000664 |  |
| **Chemical**<br>**compound,**<br>**drug** | Biocytin | ThermoFisher<br>Scientific | Cat# B1592 |  |
| **Chemical**<br>**compound,**<br>**drug** | NBQX<br>disodium salt | Tocris | Cat# 1044 |  |
| **Chemical**<br>**compound,**<br>**drug** | Tetrodotoxin<br>(TTX) | Sigma Aldrich | Cat# 554412 |  |
| **Chemical**<br>**compound,**<br>**drug** | Nimodipine | Abcam | Cat# ab120138 |  |
| **Chemical**<br>**compound,**<br>**drug** | Strychnine<br>hydrochloride | Sigma | Cat# S8753 |  |
| **Chemical**<br>**compound,**<br>**drug** | SR-95531<br>hydrobromide<br>(gabazine) | Tocris | Cat# 1262 |  |
| **Chemical compound, drug** | GYKI-53655 hydrochloride | Tocris | Cat# 2555 |  |
| **Chemical compound, drug** | (+)-MK-801 hydrogen maleate | Sigma Aldrich | Cat# M107 |  |
| **Antibody** | Anti-DsRed (Rabbit polyclonal) | Takara Bio | Cat# 632496<br>RRID: AB_10013483 | IHC (1:1000) |
| **Antibody** | Anti-mCherry (Goat polyclonal) | SICGEN | Cat# AB0040-200<br>RRID: AB_2333093 | IHC (1:2000) |
| **Antibody** | Anti-GFP (Chicken polyclonal) | Aves Labs | Cat# GFP-1020<br>RRID: AB_10000240 | IHC (1:2000) |
| **Antibody** | Anti-rabbit Cy3 (Donkey polyclonal) | Jackson ImmunoResearch Labs | Cat# 711-165-152<br>RRID: AB_2307443 | IHC (1:500) |
| **Antibody** | Anti-goat Cy3 (Donkey polyclonal) | Jackson ImmunoResearch Labs | Cat# 705-165-147<br>RRID: AB_2307351 | IHC (1:1000) |
| **Antibody** | Anti-chicken Alexa Fluor 488 (Donkey polyclonal) | Jackson ImmunoResearch Labs | Cat# 703-545-155<br>RRID: AB_2340375 | IHC (1:500) |
| **Antibody** | Streptavidin-conjugated Alexa Fluor 488 | Molecular Probes | Cat# S32354<br>RRID: AB_2315383 | IHC (1:500) |
| **Recombinant DNA reagent** | AAV1-CAG-Flex-mRuby2-GSG-P2A-GCaMP6f-WPRE-pA | addgene | Cat# 68719-AAV1 |  |
| **Recombinant DNA reagent** | AAV1-CAG-ChR2-Venus-WPRE-SV40 | addgene | Cat# 20071-AAV1 |  |
| **Recombinant DNA reagent** | AAVrg-Syn-ChR2(H134R)-GFP | addgene | Cat# 58880-AAVrg |  |
| **Software, algorithm** | pClamp 10 | Molecular Devices | RRID: SCR_011323 |  |
| **Software, algorithm** | SutterPatch | Sutter Instrument |  |  |
| **Software, algorithm** | Igor Pro 8 | Wavemetrics | RRID:SCR_000325 |  |
| **Software, algorithm** | ZEN Blue | Zeiss | RRID:SCR_013672 |  |
| **Software, algorithm** | Fiji | Fiji.sc | RRID:SCR_002285 |  |
| **Software, algorithm** | Neurolucida | MBF Bioscience | RRID: SCR_001775 |  |
| **Software, algorithm** | Excel | Microsoft | RRID:SCR_016137 |  |
| **Software, algorithm** | GraphPad Prism 9 | GraphPad | RRID:SCR_002798 |  |
| **Software, algorithm** | Affinity Designer | Serif | RRID:SCR_016952 |  |

